# A systematic evaluation of the design, orientation, and sequence context dependencies of massively parallel reporter assays

**DOI:** 10.1101/576405

**Authors:** Jason Klein, Vikram Agarwal, Fumitaka Inoue, Aidan Keith, Beth Martin, Martin Kircher, Nadav Ahituv, Jay Shendure

## Abstract

Massively parallel reporter assays (MPRAs) functionally screen thousands of sequences for regulatory activity in parallel. Although MPRAs have been applied to address diverse questions in gene regulation, there has been no systematic comparison of how differences in experimental design influence findings. Here, we screen a library of 2,440 sequences, representing candidate liver enhancers and controls, in HepG2 cells for regulatory activity using nine different approaches (including conventional episomal, STARR-seq, and lentiviral MPRA designs). We identify subtle but significant differences in the resulting measurements that correlate with epigenetic and sequence-level features. We also test this library in both orientations with respect to the promoter, validating *en masse* that enhancer activity is robustly independent of orientation. Finally, we develop and apply a novel method to assemble and functionally test libraries of the same putative enhancers as 192-mers, 354-mers, and 678-mers, and observe surprisingly large differences in functional activity. This work provides a framework for the experimental design of high-throughput reporter assays, suggesting that the extended sequence context of tested elements, and to a lesser degree the precise assay, influence MPRA results.

## INTRODUCTION

The spatiotemporal control of gene expression is orchestrated in part by distally located DNA sequences known as enhancers. The first viral and cellular enhancers were identified by cloning fragments of DNA into a plasmid with a reporter gene and promoter^1–4^. At present, enhancement of transcription in such a reporter assay remains the common standard for evaluating whether a putative regulatory element is a *bona fide* enhancer. However, conventional, one-at-a-time reporter assays are insufficiently scalable to test the >1 million putative enhancers in the human genome^5–8^.

Massively parallel reporter assays (MPRAs) modify the *in vitro* reporter assays described above to facilitate simultaneous testing of thousands of putative regulatory elements or variants thereof^9–11^ in a single experiment. Instead of relying on measurement of a conventional reporter, MPRAs characterize each element in a multiplex fashion, through sequence-based quantification of barcodes incorporated into the RNA, each associated with a different element^10^. MPRAs (a term we use broadly to encompass related methods including STARR-seq^12^ and lentiMPRAs^13^) have facilitated the scalable study of putative regulatory elements for goals ranging from functional annotation^12–14^ to variant effect prediction^10, 11, 15–18^ to evolutionary reconstruction^19, 20^.

To date, several groups have implemented enhancer-focused MPRAs, but with diverse designs. Some of the major differences include whether the enhancer is upstream^10, 11^ vs. within the 3’ UTR of the reporter gene^12^, and whether the construct remains episomal vs. integrated^13^. Additionally, MPRAs test sequences cloned in one of two possible orientations, effectively assuming that enhancer activity is independent of orientation. Finally, while larger sheared genomic DNA fragments^12^, PCR amplicons^15^ or captured sequences^21, 22^ have been used in MPRAs, most studies using MPRAs synthesize libraries of putative enhancers on microarrays, and are therefore limited to testing shorter sequences (typically less than 200 bp).

Unfortunately, we have, as a field to date, largely failed to evaluate how these design choices impact or bias the results of MPRAs. First, although assays like STARR-seq wherein the enhancer serves as the barcode are more straightforward to implement, it is unknown how the position (3’ of the promoter, rather than 5’ as in a more conventional reporter assay vector) or the fact that the sequence is serving as the barcode, influences results. Second, although we previously showed substantial differences between episomal vs. integrated MPRAs^13^, it is not clear how these differences rank relative to other design choices. Third, although the orientation-independence of enhancers has been shown in a few cases, to our knowledge the robustness of this assumption has never been tested in a systematic fashion with a large number of different sequences. Finally, the typical choice to test <200 bp fragments, each corresponding to a putative enhancer, is entirely based on technical limitations of massively parallel DNA synthesis, rather than on any principled understanding of the actual size of enhancers. The consequences of this choice for the results obtained remain largely unquantified.

Particularly as efforts to validate the >1 million putative human enhancers^5–8^, as well as the growing number of disease-associated noncoding variants, begin to scale, a clear-eyed understanding of the biases and tradeoffs introduced by various MPRA experimental design choices is needed. To this end, we performed a systematic comparison, testing the same 2,440 sequences for regulatory activity using nine different MPRA strategies, including conventional episomal, STARR-seq, and lentiviral designs. Second, we tested the same sequences in both orientations relative to the promoter. Finally, we further developed a multiplex pairwise assembly protocol^23^, and applied it to test short (192 bp), medium (354 bp), and long (678 bp) versions of the same enhancers. Our results quantify the impact of MPRA experimental design choices and also provide further insight into the nature of enhancers.

## RESULTS

### Implementation and testing of nine distinct MPRA strategies

We sought to systematically compare nine different MPRA strategies. They are as follows, with the numbers corresponding to the vertical order on the left side of **Figure 1**: (**1**) the “classic” MPRA assay design, using the pGL4.23c vector, wherein the enhancer library resides upstream of a minimal promoter, and the associated barcodes reside in the 3’ UTR of the reporter gene (“pGL4”)^10, 24^. (**2-3**) the Self-Transcribing Active Regulatory Region Sequencing (STARR-seq) assay design, wherein the enhancer library resides in the 3’ UTR of the reporter gene, either as originally described (“HSS”)^12^ or a newer version that uses the bacterial origin of replication for transcription initiation (“ORI”)^25^. In both cases, we introduce barcodes immediately adjacent to the location of the enhancers in the 3’ UTR in order to facilitate consistent procedures with other assays. (**4, 6, 8**): The lentiMPRA assay design, wherein lentiviral integration of a library of reporters is used to mitigate the concern that episomes lack native chromatin, either with the enhancer library upstream of the minimal promoter and the associated barcodes in the 3’ UTR of the reporter gene (5’/3’ WT)^13^, the enhancer library upstream of the minimal promoter and the associated barcodes in the 5’ UTR of the reporter gene (5’/5’ WT), or with both the enhancer library and the associated barcodes in the 3’ UTR of the reporter gene (3’/3’ WT). The 5’/5’ WT design was developed in light of evidence for distance-dependent template switching prior to lentiviral integration^26, 27^, as it reduces the distance between the enhancer and barcode from 801 bp (5’/3’ WT) to 102 bp (5’/5’ WT). The 3’/3’ WT design is analogous to STARR-seq, but integrated into the genome, and also addresses template switching by positioning the enhancer and barcode immediately adjacent to one another in the 3’ UTR. (**5, 7, 9**): These designs are identical to the three lentiMPRA designs, except that the vector harbors a mutant integrase such that the constructs remain episomal (5’/3’ MT, 5’/5’ MT, 3’/3’ MT)^13^.

**Figure 1.**
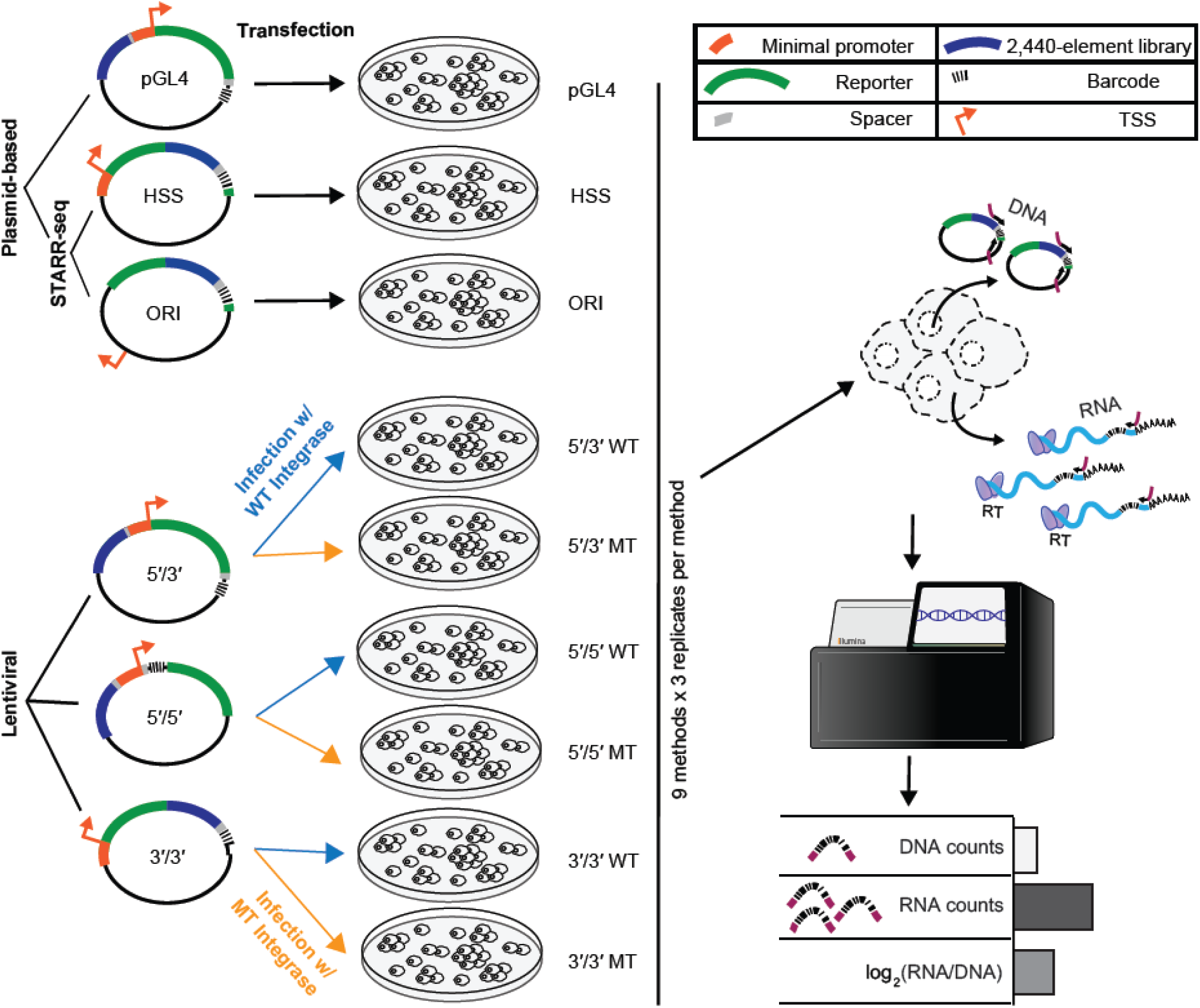
Nine MPRA strategies and experimental workflow. Nine different MPRA designs were tested. These are schematically represented on the left, and from top to bottom include pGL4.23c (pGL4); the original STARR-seq vector (HSS); STARR-seq with no minimal promoter (ORI); and lentiMPRAs with the enhancer library upstream of the minimal promoter and the associated barcodes in the 3’ UTR of the reporter gene (5’/3’), the enhancer library upstream of the minimal promoter and the associated barcodes in the 5’ UTR of the reporter gene (5’/5’), or with both the enhancer library and the associated barcodes in the 3’ UTR of the reporter gene (3’/3’). The episomal designs (pGL4, HSS, ORI) were transfected into HepG2 cells, while 5’/5’, 5’/3’, and 3’/3’ were packaged with either wild type (WT) or mutant (MT) integrase and infected into HepG2 cells. DNA and RNA were extracted from the cells, and the enhancer-associated barcodes amplified and sequenced. Finally, the activity score of each element was computed as the log_2_ of the normalized count of RNA molecules from all barcodes corresponding to the element, divided by the normalized number of DNA molecules from all barcodes corresponding to the element.

For a common set of sequences that could be tested in conjunction with each of these nine MPRA strategies, we turned to a library that we previously described when developing lentiMPRAs^13^. Briefly, this library consists of 2,236 candidate enhancer sequences based on ChIP-seq peaks in the HepG2 cell line, along with 204 controls (**Supplemental Table 1**). Of note, our criteria did not require that candidate enhancers fall distal of promoters, and as such 281 of the 2,236 overlap promoters (defined as +/− 1 kilobase (Kb) of the TSS of a protein-coding gene). The controls consist of synthetically designed sequences that previously demonstrated enhancer MPRA activity (100 positive controls) or lack thereof (100 negative controls)^28^ in HepG2 cells, along with 2 positive and 2 negative controls derived from endogenous sequences that we validated with luciferase assays^13^. All sequences were 171 base pairs (bp) in length and were synthesized on a microarray together with common flanking sequences (total length of 230 bp). A 15 bp degenerate barcode was appended during library amplification, and the amplicons cloned to the human STARR-seq (HSS) vector. The enhancer/barcode region of the HSS library was amplified and used for two purposes – first, it was sequenced in order to determine which barcodes were associated with which enhancers; second, the amplicons were cloned to other vectors to create libraries for the remaining eight MPRA designs (**Supplemental Figure 1**). As such, the relative abundances of enhancers and barcodes, as well as the enhancer-barcode associations, were consistent across all MPRA libraries. Cloning details and references for each of the nine assay designs are provided in the **Methods**.

Plasmid libraries were transfected into HepG2 cells in triplicate (three different days), and lentiMPRA libraries were packaged with either wild-type (WT) or defective (MT) integrase lentivirus and infected into HepG2 in triplicate (three different days). We extracted DNA and RNA from cells, amplified barcodes via PCR and RT-PCR respectively, and sequenced the amplicons to generate barcode counts (**Figure 1**). An activity score for each element was calculated as the log_2_ of the normalized count of RNA molecules from all barcodes corresponding to the element, divided by the normalized number of DNA molecules from all barcodes corresponding to the element (**Supplemental Table 2**). For each of the 27 experiments (9 assays × 3 replicates), only barcodes observed in both RNA and DNA were considered. For 26 of 27 experiments (all but 3’/3’ MT replicate 1), the median number of barcode counts per element was greater than 100 (**Supplemental Figure 2**).

### Different MPRA designs yield different results

We first sought to evaluate the technical reproducibility of each assay. Most of the assays were highly correlated between the three replicates. Specifically, the intra-assay Pearson correlations for pairwise comparisons of activity scores of replicates exceeded 0.90 for all assays except for the 5’/3’ MT (mean *r* = 0.87) and 3’/3’ MT (mean *r* = 0.54) assays (**Figure 2A**; **Supplemental Figure 3A**). Similarly, we confirmed reasonable correlation for 5’/3’ WT and 5’/3’ MT between this and our previous study^13^ (*r* = 0.92 for 5’/3’ WT and *r* = 0.81 for 5’/3’ MT, **Supplemental Figure 3B**).

**Figure 2.**
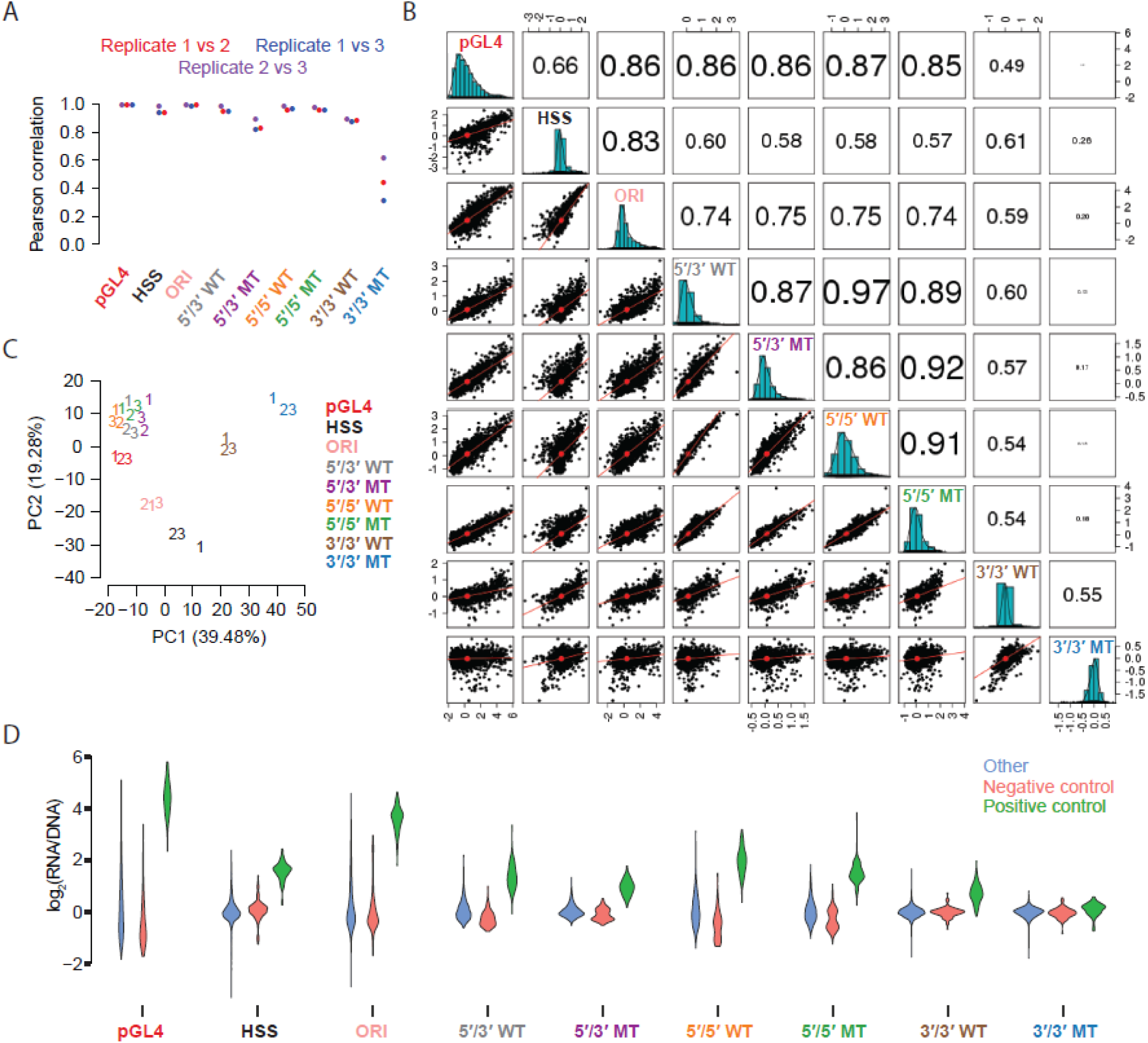
Quantitative comparison of different MPRA strategies. **(A)** Beeswarm plot of the Pearson correlation values for each of the three possible pairwise comparisons among the replicates of each MPRA technique. **B)** Scatter matrix displaying scatter plots corresponding to each of the 36 pairs of possible inter-assay comparisons (lower diagonal elements). Shown on the diagonal is a histogram of the log_2_(RNA/DNA) ratios, averaged among replicate samples. Also shown are Pearson correlation values among each pair of comparisons, with the size of the text proportional to the magnitude of the correlation coefficient (upper diagonal elements). See also **Supplemental Figure 5** for the equivalent result but with Spearman correlations. **C)** Principal components analysis (PCA) of 27 experiments, *i.e.* three replicates (labeled ‘1’, ‘2’, and ‘3’) of each of the nine different MPRA approaches tested. Shown are the first two principal components that together explain over half of the variation. Slight jitter was added to each data point to enhance readability. **D)** Violin plots displaying the distribution of average log_2_(RNA/DNA) ratios across independent transfections for positive controls, negative controls, and putative enhancer sequences tested, for each of the nine assays.

We next sought to compare the results of the various assay designs to one another. To do this, we calculated the average activity score for each element across all technical replicates of a given assay (**Supplemental Table 2**), and then compared the assays to one another. Six of the nine assays demonstrated consistently reasonable inter-assay Pearson and Spearman correlations of >0.7 with all other members of this group (**Figure 2B, Supplemental Figure 4, Supplemental Figure 5**). These were the ORI and pGL4, together with both versions of the 5’/5’ and 5’/3’ assays (*i.e.* WT and MT). The remaining three assays did not show good agreement with the other six assays, nor with one another. These three were 3’/3’ MT, which as mentioned above exhibited poor technical reproducibility, as well as 3’/3’ WT and HSS.

As a different approach to compare the assays, we subjected activity scores from all 27 experiments (9 assays × 3 replicates) to Principal Components Analysis (PCA) (**Figure 2C**). The aforementioned six assays with inter-assay correlations of >0.7 clustered reasonably closely. Interestingly, Principal Component 1 tended to separate the assays wherein the enhancer resides upstream of the minimal promoter (5’/5’, 5’/3’, and pGL4) from the assays where it resides 3’ of the reporter gene (3’/3’, HSS, and ORI). In contrast, Principal Component 2 tended to separate lentiviral designs (5’/5’, 5’/3’, and 3’/3’) from plasmid-based designs (pGL4, HSS, and ORI). This suggests systematic differences in the enhancer activity measurements that relate to aspects of MPRA design.

Next, we examined the dynamic range of activity scores with each of the nine assays (**Figure 2D**). The classic enhancer reporter vector (pGL4) and the promoter-less STARR-seq assay (ORI) exhibited the greatest dynamic range, with pGL4 showing the largest separation between the positive and negative controls (two-sided *t*-statistic = 37.46). Among the lentiviral assays, the 5’/5’ WT design exhibited the greatest dynamic range and separation of controls (two-sided *t*-statistic = 30.92).

We next sought to ask whether we could predict enhancer activity, as measured by the different assays^13^. We generated lasso regression models based on a set of 915 biochemical, evolutionary, and sequence-derived features (**Supplemental Table 3**, **Supplemental Table 4**) and a 10-fold cross-validation (CV) scheme. We were able to best predict enhancer activities for the six aforementioned assays that exhibited reasonable inter-assay consistency (Pearson’s *r* ranging from 0.59 for 5’/3’ WT to 0.71 for pGL4) (**Figure 3A**, **Supplemental Figure 6**).

**Figure 3.**
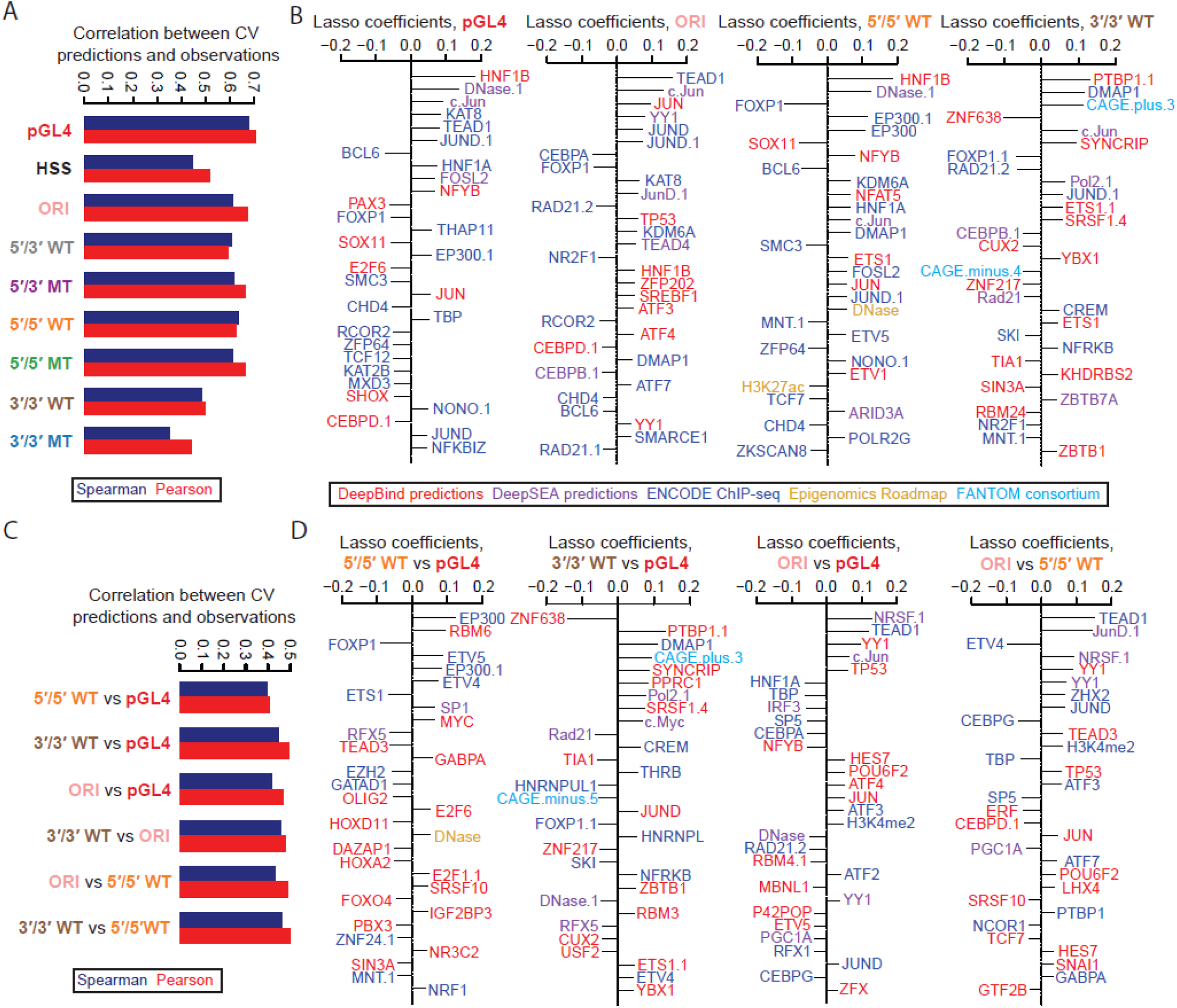
Predictive modeling of the ratios and differences between MPRA methods. **A)** Pearson and Spearman correlation coefficients for 10-fold cross-validated predictions vs. observed values for each of the nine methods. **B)** The top 30 coefficients derived from a lasso regression model trained on the full dataset derived from the pGL4 (left panel), ORI (middle-left panel), 5’/5’ WT (middle-right panel), and 3’/3’ WT (right panel) methods. Features with the extension “.1”, “.2”, etc allude to redundant features or replicate samples. See **Supplemental Figure 7** for the top 30 coefficients for the remaining five methods. **C-D)** These panels are similar to panels A and B, respectively, but show results for regression models predicting observed differences for the six possible pairwise comparisons of four of the methods. See **Supplemental Figure 8** for the top model coefficients of the remaining two pairwise comparisons of methods not shown in panel D.

Many of the top coefficients fit by these models correspond to ChIP-seq signal or sequence-based binding site predictions for transcriptional activators (*e.g.* Jun, HNF1, FOSL2, TEAD1, YY1), coactivators (*e.g.* p300, KDM6A), and repressors (FOXP1, RCOR2, HDAC2/6, EZH2, and cohesin-complex members such as SMC3 and RAD21) (**Figure 3B**, **Supplemental Figure 7**, **Supplemental Table 5**). We caution that the interpretation of feature selection and coefficient-based ranking is inherently limited by the substantial degree of multicollinearity among features (**Supplemental Table 4**), which in turn limits the determination of which features are mechanistically or causally involved. Potential reasons for inter-feature correlations include: i) several paralogous factors may bind to nearly identical motifs, leading to overlapping sequence-based binding predictions, ii) several factors may bind in a complex, leading to a redundant ChIP-seq signal, and iii) an upstream factor might lead to the deposition or removal of multiple epigenetic marks simultaneously, leading to a correlated or anti-correlated signal. Of note, another potential confounder is that the criteria used to select elements relied on features including FOXA1/2, HNF4A, EP300, H3K27ac, CHD2, RAD21, and SMC3 ChIP peaks^13^, which could exaggerate the contributions of these factors, or bias the contributions of other features, in models attempting to predict general enhancer activity in HepG2 cells.

We next sought to ask whether we could predict differences in enhancer activity between the assays, based on the same set of 915 features. For models predicting pairwise differences between the results of the pGL4, 5’/5’ WT, 3’/3’ WT, and ORI assays, we were able to achieve correlations between 0.4-0.5 (**Figure 3C**, **Supplemental Figure 8A**). We were particularly interested in whether features corresponding to RNA binding proteins and splicing factors would be especially predictive of the promoterless STARR-seq (ORI) or 3’/3’ WT results, as in these assays the enhancer itself is included in the 3’ UTR of the reporter. Indeed, SRSF1/2, BRUNOL4, PTBP1, PPRC1, KHDRBS2, SYNCRIP, and MBNL1, which are known to modulate mRNA stability and splicing, predict differences in measured activity in ORI or 3’/3’ WT vs. 5’/5’ WT or pGL4 (**Figure 3D**, **Supplemental Figure 8B**, **Supplemental Table 5**). Several promoter and enhancer binding proteins such as NRSF1, TEAD1, Jun, YY1, ETV4/5, TBP, CEBPG, CEBPD, and CEBPA are also predictive of differences, both between ORI and the other assays as well as between 5’/5’ WT vs. pGL4, suggesting that to some degree, these factors act in a context-dependent manner. Interestingly, general transcriptional activity, as measured by CAGE, was among the most predictive features of the 3’/3’ WT context (**Figure 3D**, **Supplemental Figure 8B**). As this is the only assay where the tested elements are both genomically integrated and distally located from the promoter, this observation suggests CAGE-based transcriptional activity may be a good predictor of distal enhancer activity^29, 30^.

### Enhancer activity is largely, but not completely, independent of sequence orientation

We next set out to test a key aspect of the canonical definition of enhancers, that they function independently of their orientation with respect to the promoter. To assess whether orientation influences enhancer activity in MPRAs, we directionally cloned 2,336 sequences (the same 2,236 candidates described above extended out to 192 bp, along with 50 positive and 50 negative controls from Vockley *et al.*^15^), in both orientations into the pGL4 vector and pooled these two libraries together (**Figure 4A**). We transfected our pGL4 dual-orientation library into HepG2 cells in quadruplicate. The median number of barcode counts per element was greater than 100 (**Supplemental Figure 9**), and the measured activities were highly correlated between replicates (Pearson’s *r* > 0.98; **Figure 4B**, **Supplemental Figure 10**, **Supplemental Table 6**). Interestingly, enhancer activities for the same elements cloned in forward vs. reverse orientation to the pGL4 vector were also highly correlated in pairwise comparisons between replicates (mean *r* = 0.88), but consistently less so than same-orientation comparisons (*r* > 0.98; **Figure 4B**). This suggests that the activity of enhancers, is largely, but not completely, independent of orientation.

**Figure 4.**
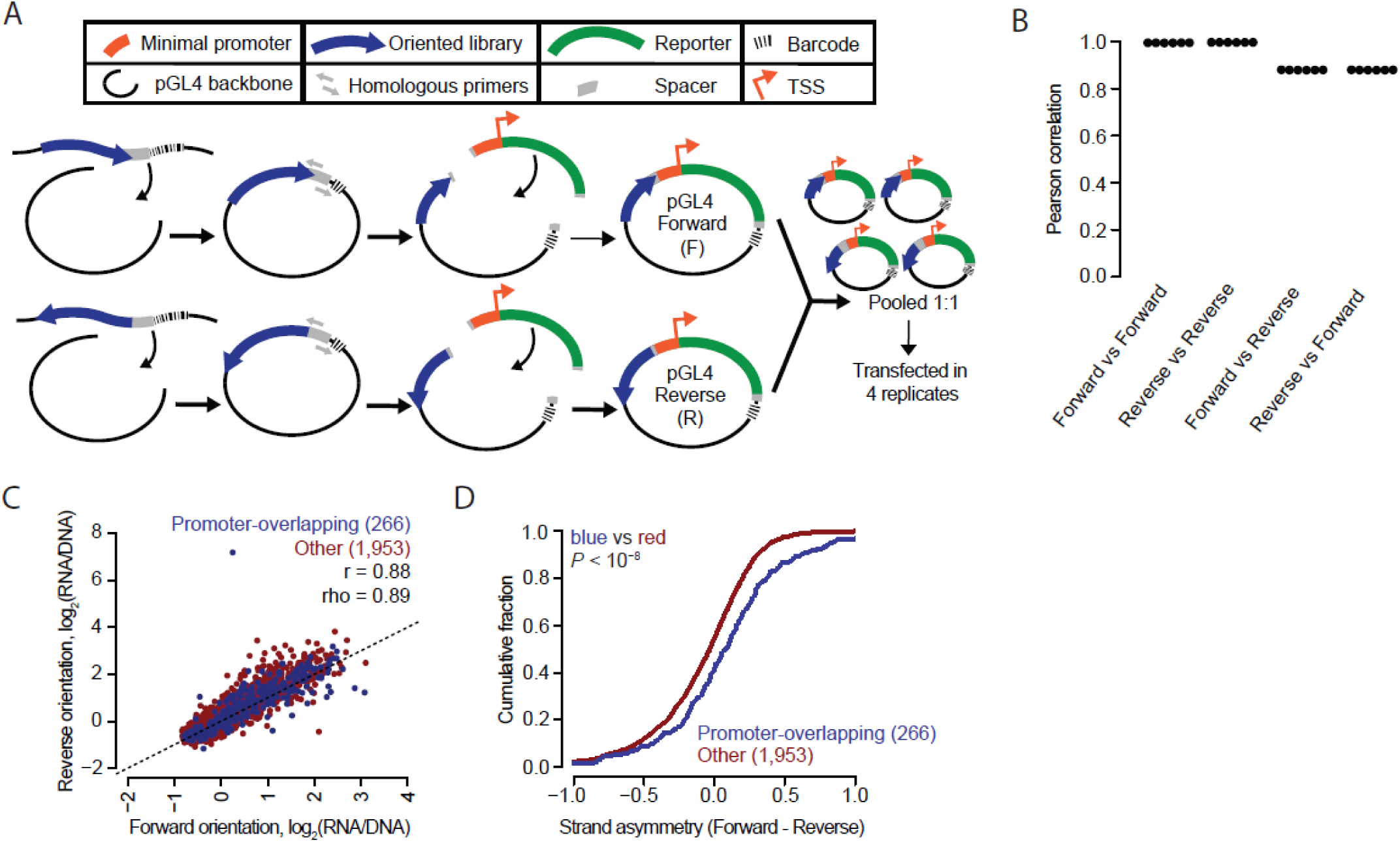
Enhancer activity is largely, but not completely, independent of sequence orientation. **A)** A schematic of the experimental workflow used to produce an MPRA library with each element in both orientations. The 2,336 element library was cloned into the pGL4 backbone in both orientations as two separate libraries. These were then pooled together in equal amounts and transfected into HepG2 cells in quadruplicate. **B)** Beeswarm plot of the Pearson correlation values corresponding to each of the six possible pairwise comparisons among the four replicates. The correlations are computed between observed enhancer activity values for elements positioned either in the same (Forward vs. Forward and Reverse vs. Reverse) or opposite (Forward vs. Reverse and Reverse vs. Forward) orientations. **C)** Scatter plots of the average activity score of each element in the Forward vs. Reverse orientation, split out by promoter-overlapping (blue; +/− 1 Kb of the TSS of a protein-coding gene) and other (red) elements. **D)** Cumulative distributions measuring strand asymmetry between promoter-overlapping elements and other elements. Here, “Forward” and “Reverse” were defined as “sense” and “antisense”, respectively, in relation to the orientation of the TSS for promoter-overlapping elements (n = 266); and were defined as “plus” and “minus”-stranded, respectively, in relation to the chromosome annotation for other elements (n = 1,953). Similarity of the blue distribution to that of the red was tested (one-sided Kolmogorov–Smirnov [K–S] test, *P* value).

In contrast with enhancers, promoters are established to be directional in activity^31, 32^. 266 of 281 promoter-overlapping elements were successfully measured in both orientations. We therefore tested whether these behaved differently than 1,953 more distally located elements. Indeed, the promoter-overlapping sequences exhibited greater differences in activity between the two orientations than the distal elements, supporting the conclusion that they inherently contain signals to promote transcription in an asymmetric fashion relative to enhancers (**Figure 4C-D**).

### Including additional sequence context around tested elements leads to significant differences in the results of MPRAs

Most MPRAs use array-synthesized libraries that are limited in length, typically to less than 200 bp in terms of the assayed sequence. The impact of this length restriction, which is technically rather than biologically motivated, has not been rigorously explored. To evaluate this, we designed 192 bp (“short”), 354 bp (“medium”), and 678 bp (“long”) versions of our candidate enhancer library, centered at the same genomic position, and corresponding to the equivalent 2,236 candidate enhancers tested above (*i.e.*, for longer versions, simply including more flanking sequence from the reference genome; **Supplemental Table 1**). We also included 50 high and low-scoring putative elements from Vockley *et al.*^15^ in the short and medium libraries.

The 192 bp versions of these candidate enhancers, which were utilized for the orientation experiments above, were synthesized directly on a microarray as before; sequencing showed a 100% yield (2,336/2,336) and a 3.8-fold interquartile range (IQR) for relative abundance (**Supplemental Figure 11A**). To generate the 354 bp versions, we performed Multiplex Pairwise Assembly (MPA)^23^ on overlapping pairs of array-synthesized 192 bp fragments; sequencing showed a 95% yield (2,241/2,336) and a 4.9-fold IQR (**Supplemental Figure 11A**). Finally, to generate the 678 bp versions, we developed a “two-round” version of MPA, which we call Hierarchical Multiplex Pairwise Assembly (HMPA; **Supplemental Figure 11B**, **Supplemental Figure 12**). HMPA of overlapping pairs of array-synthesized 192 bp fragments yielded overlapping pairs of 354 bp fragments, which were further assembled to generate 678 bp fragments. These 678 bp fragments had an 84% yield (1,887/2,236) and 27.9-fold IQR (**Supplemental Figure 11A**). We verified a subset of our long enhancers with PacBio sequencing (**Supplemental Figure 11C-D**; chimera rate of 16.5%).

We cloned all three libraries into the pGL4 vector, pooled them together, and transfected this pool in quadruplicate to HepG2 cells (**Figure 5A**). We sequenced barcodes in RNA and DNA and calculated activity scores as above (**Supplemental Table 7**). Requiring each element to be detected with at least 10 unique barcodes, there were 2,109 candidate enhancers tested at both short and medium lengths, 658 tested at both medium and long lengths, and 670 tested at both short and long lengths. The median number of barcode counts per element was greater than 100 for all replicates (**Supplemental Figure 13**). Technical replicates within any given length class were highly reproducible, albeit modestly less so for long elements (mean Pearson’s *r* = 0.94, **Figure 5B**, **Supplemental Figure 14**). However, there was substantially less agreement for the same candidate enhancers tested at different lengths, with correlations appearing to decrease as a function of the length discrepancy between tested elements (short vs. medium, mean *r* = 0.78; medium vs. long, mean *r* = 0.67; short vs. long, mean *r* = 0.53; **Figure 5B-C**). Finally, we observed that the positive control sequences were significantly more active than the negative controls when tested as either 192 bp or 354 bp fragments (*P* < 0.01, Wilcoxon sign-rank test; **Figure 5D**; of note, the positive controls were only included in the short and medium libraries, such that equivalent comparisons for the long libraries cannot be performed).

**Figure 5.**
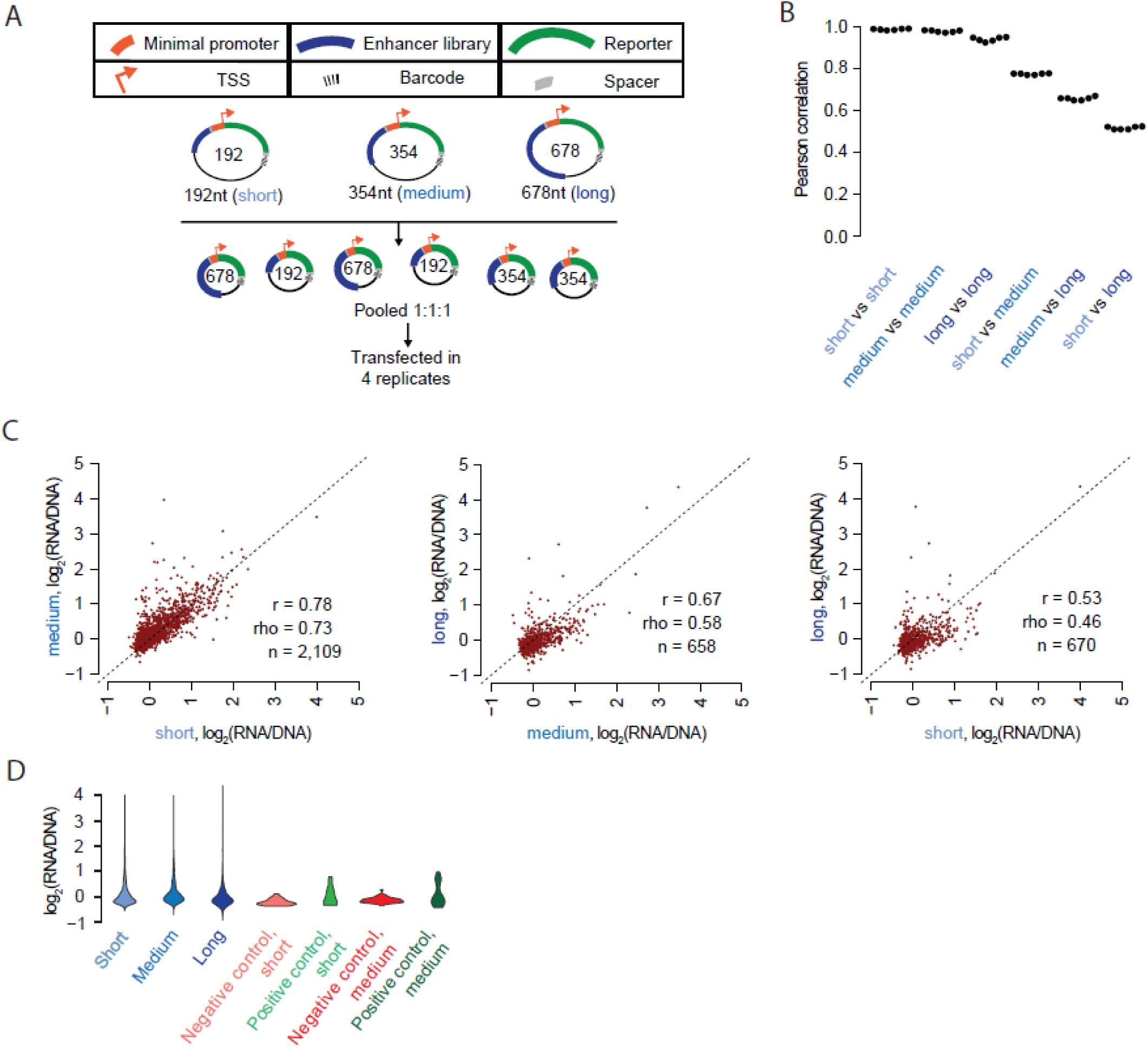
Including additional sequence context around tested elements leads to significant differences in the results of MPRAs. **A)** Experimental schematic. 192 bp, 354 bp, and 678 bp libraries were synthesized, assembled, and cloned into the pGL4 backbone. These were pooled and transfected into HepG2 cells in quadruplicate. **B)** Beeswarm plot of the Pearson correlation values corresponding to each of the six possible pairwise comparisons among the four replicates. The correlations are computed between observed enhancer activity values for elements measured in each of the three possible size classes. **C)** Scatter plots of the average activity score of each element, comparing short vs. medium, medium vs. long, and short vs. long versions of each element. **D)** Violin plot displaying the distribution of average log_2_(RNA/DNA) ratios for short, medium, and long versions of the elements tested, as well as for positive and negative controls at short and medium lengths.

We trained lasso regression models to predict activities using features which were re-computed for each of the three size classes (**Figure 6A**, **Supplemental Figure 15**, **Supplemental Table 4**). The lower performance of the model for long element library is possibly consequent to its fewer sequences, lower technical reproducibility, or an increase in the effect of non-linear interactions between features that reduce predictive performance. Known predictors of enhancer activity were consistently present in the top coefficients, although their relative rankings differed depending on the size class being examined (**Figure 6B**, **Supplemental Table 5**). Next, we sought to explicitly model how *differences* in predicted factor binding might explain *differences* in enhancer activity, as measured by different pairs of size classes. For example, in attempting to explain observed activity differences in long vs. short elements, we computed a set of features as the differences in predicted binding, or measured ChIP-seq signal, between the long element and corresponding short element (e.g., ΔARID3A = ARID3A_long_ – ARID3A_short_). Many of the top features originated from sequence-based differences in predicted binding in the extra genomic context surrounding the core element. Features consistently observed to explain activity differences in longer elements include RPC155, the catalytic core and largest component of RNA polymerase III; Jun and FOS, components of the AP-1 complex; ATF2, EZH2, and HDAC1/2, core histone-modifying enzymes; and the transcription factors ARID3A, DRAP1, and SP1/2/3 (**Figure 6C-D**, **Supplemental Table 5**).

**Figure 6.**
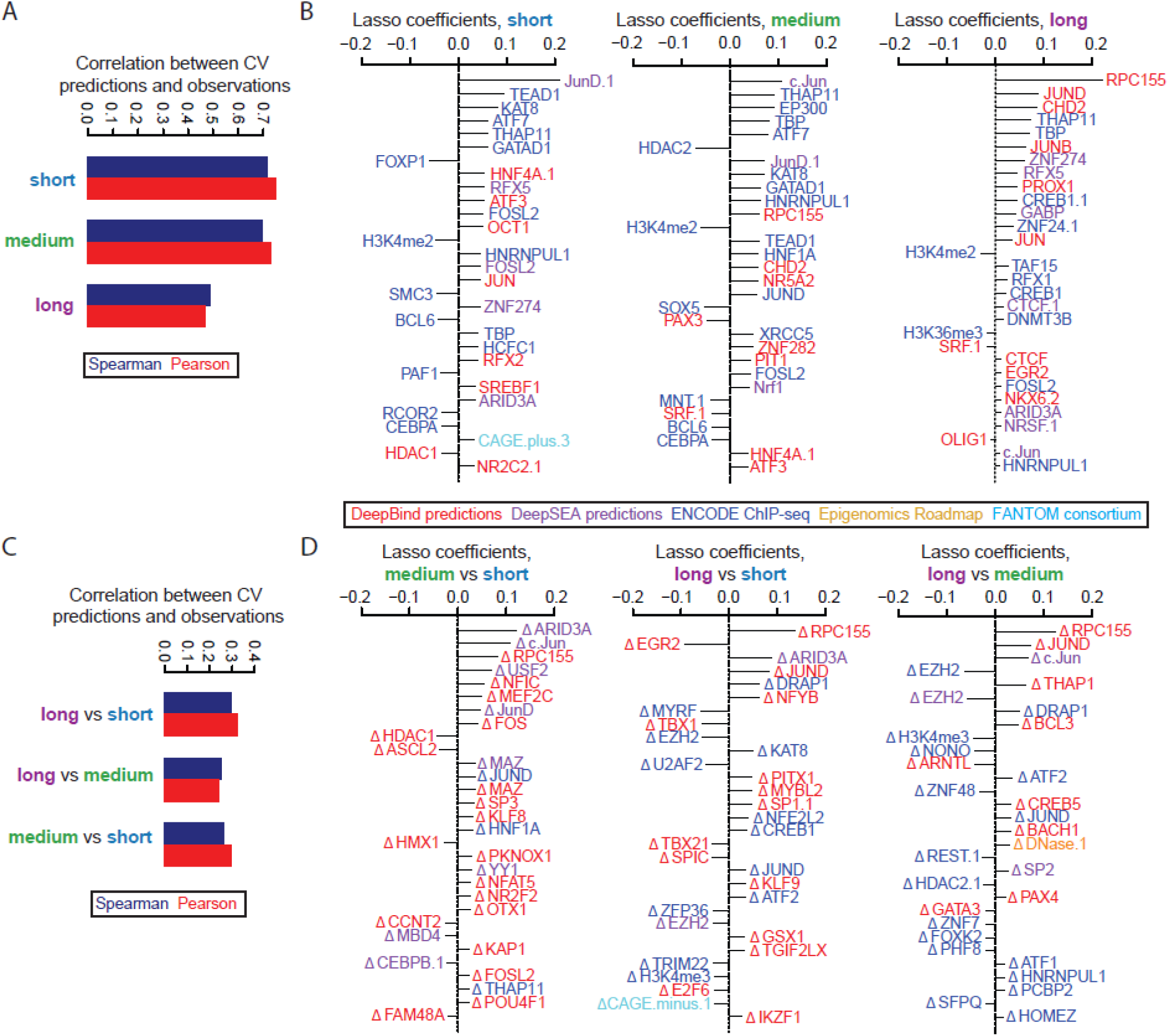
Predictive modeling of factors dependent on element size. **A)** Pearson and Spearman values between the 10-fold cross-validated predictions and observed values for each of the three size classes tested. **B)** The top 30 coefficients derived from a lasso regression model trained on the full dataset derived from the short (left panel), medium (middle panel), and long (right panel) size classes. Features with the extension “.1”, “.2”, etc allude to redundant features or replicate samples. **C-D)** These panels are similar to panels A and B, respectively, but show results for regression models attempting to predict observed differences in activity for the same elements tested at different lengths, using features that quantify the differences in signal between the longer and shorter versions of the elements.

## DISCUSSION

Over the past decade, MPRAs have enabled researchers to functionally test large numbers of DNA sequences for regulatory activity and in the process address numerous biological questions. While different groups utilize various backbones and assay designs, there has been no systematic comparison of how these different strategies influence results.

Here, we have sought to perform a systematic comparison of all major MPRA strategies, and to concurrently investigate the consequences of other key design choices, *i.e.* element orientation and element length. We generally observe concordance between different MPRA designs, albeit to varying degrees. Six of the nine assays exhibited both technical reproducibility as well as reasonable agreement with one another (pGL4, ORI, 5’/5’ WT, 5’/5’ MT, 5’/3’ WT, 5’/3’ MT). Furthermore, as we previously showed for the 5’/3’ WT and 5’/3’ MT assays, enhancer activities as measured by MPRAs^13^ are reasonably well predicted by models based on primary sequence together with biochemical measurements at the corresponding genomic locations. Nonetheless, despite the general agreement, there are clear systematic differences that are predictable by similar features, consistent with what we previously observed in comparing 5’/3’ WT vs. 5’/3’ MT MPRAs^13^. Taken together, our results support a view wherein diverse MPRAs are all measuring enhancer activity, but design differences (*e.g.* integrated vs. episomal; 5’ vs. 3’ location of the enhancer) influence the results to a modest degree. For example, features influencing mRNA stability and splicing predict differences between the pGL4 and 5’/5’ WT assays (enhancer upstream of promoter) vs. the ORI and 3’/3’ WT assays (enhancer in 3’ UTR and transcribed).

Overall, our results support show a preference for three of the nine MPRA designs evaluated here—pGL4, ORI, and 5’/5’ WT, which all had reasonable inter-assay correlations. The pGL4 assay has the advantage of representing the “classic” enhancer reporter assay design, had the greatest dynamic range, and was the most predictable with our lasso regression, but the disadvantage of being episomal rather than integrated. The ORI assay (*i.e.* promoterless STARR-seq) has the advantage of eliminating the need to associate barcodes, potentially allowing for greater library complexities, and has a large dynamic range, but the disadvantages of confounding enhancer activity with possible effects on mRNA splicing and/or stability, and also of being episomal rather than integrated. The 5’/5’ WT assay has the advantage of being integrated rather than episomal, and—amongst lentiviral assays—mitigates the template switching issue by minimizing the distance between the enhancer and barcode. However, template switching still occurs to some degree, and the assay exhibits a lower dynamic range than pGL4 or ORI assays.

Another key finding is our confirmation that the activity of enhancers, is largely, but not completely, independent of orientation, at least as measured by MPRAs. This is of course part of the original definition of enhancers^1^, but efforts to systematically test the validity of this assumption across a large number of sequences have been limited. Candidate enhancer sequences derived from the vicinity of TSSs exhibited greater directionality, consistent with a subset of these bearing features of oriented promoters.

Finally, we developed improved methods to efficiently assemble longer DNA fragments from array-synthesized oligonucleotides, and applied them to evaluate the extent to which including additional sequence context around tested elements impacts MPRA results. We successfully assembled 95% of 2,336 × 354 bp targets using MPA, compared to just 71% of 2,271 × 192-252 bp targets in our original description of the method^23^. Moreover, our new hierarchical MPA (HMPA) is to our knowledge the first protocol to *in vitro* assemble thousands of sequences, each over 600 bp, as a single library. Unlike potential alternatives, it does not require specialized equipment, making it more widely accessible^33^.

The sub-200 bp length of subsequences typically tested is a choice related to the technical limits of microarray-based synthesis. In the genome, there are no such limits, and it remains unclear what the appropriate “enhancer size” is to test in MPRAs and whether this choice matters. To evaluate this, we tested candidate enhancers at three different lengths (192, 354 and 678 bp). We observe correlations between the same elements tested at all lengths, but these correlations clearly drop off as a function of length differences. At the extreme, the activities of 678 bp vs. 192 bp versions of the same candidate enhancers were more poorly correlated than nearly all of our inter-assay comparisons (Pearson *r* = 0.53, Spearman rho = 0.46). Furthermore, these data suggest that the longer sequences are adding biologically relevant signal, as features corresponding to relevant transcription factors explain differences in activity of longer vs. shorter sequences. For example, a feature corresponding to RPC155, the catalytic subunit of RNA polymerase III, is the strongest coefficient separating the 678 bp constructs from the 192 and 354 bp constructs, and also one of the stronger coefficients separating the 354 bp from 192 bp constructs.

In conclusion, our results favor the use of one of three MPRA designs: pGL4, ORI, and 5’/5’ WT. Each of these assays exhibited strong technical reproducibility, reasonable interassay correlation with one another despite design differences, and reasonable predictability based on sequence and biochemical features of the corresponding genomic regions. Our results also suggest a degree of caution in interpreting the results of all MPRAs, as they are all subject to influence by aspects of the assay design. Finally, we conclude that whereas MPRA results are largely independent of the orientation of tested elements, they are surprisingly dependent on the length of the elements tested—in other words, sequence context matters. Further work is necessary to develop or improve methods like HMPA to facilitate the construction of complex, uniform MPRA libraries of longer sequences, as well as to further explore the optimal parameters of element selection (*e.g.* length, centering), particularly as MPRAs are further scaled to functionally test candidate enhancers on a genome-wide basis.

## METHODS

### Design, barcoding, and cloning of the enhancer library into the human STARR-seq (HSS) vector

We used an existing array library from Inoue *et al.*^13^. This library consists of 2,440 unique 171 bp candidate enhancer sequences, based on ChIP-seq peaks in HepG2. Each sequence was flanked with a 15 bp sequence on the 5’ end (Original_Array_5adapter) and a 44 bp sequence on the 3’ end (Original_Array_3adapter) (**Supplemental Table 8**). More detail on enhancer design can be found in that manuscript^13^. We first amplified the library using the following primers: STARR-Seq-AG-f and spacer-AG-r (**Supplemental Table 8**). These amplify the library excluding the previously designed barcodes, while adding homology to the STARR-seq vector (Addgene ID:71509)^12^ on the 5’ end and a spacer sequence on the 3’ end that we use for all subsequent libraries. We amplified 10 ng of array oligos with KAPA HiFi 2x Readymix (Kapa Biosystems) with a 65 °C annealing temperature and 30 s extension, following the manufacturer’s protocol. We followed the reaction in real time using Sybr Green (Thermo fisher scientific), and stopped the reaction before plateauing, after 10 cycles. We then purified the PCR product with a 1.8x AMPure XP (Beckman coulter) cleanup and eluted in 50 µl of Qiagen Elution Buffer (EB), following the manufacturer’s protocol. We took 1 µL of purified PCR product and amplified in triplicate a second reaction using Kapa HiFi 2x Readymix using primers STARR-Seq-AG-f and STARR-BC-spacer-r with a 35 s extension time and 65 °C annealing temperature for eight cycles (**Supplemental Table 8**). This round of PCR added a 15 bp degenerate barcode on the 3’ end of the spacer as well as homology arms to the 3’ end of the human STARR-seq vector. We then pooled the three reactions together, ran on a 1.5% agarose gel, and gel extracted the amplicon using the QIAquick Gel Extraction Kit (Qiagen), following the manufacturer’s protocol, eluting in 17.5 µL of Qiagen EB. We then cloned a 2:1 molar excess of our gel-extracted insert into 100 ng of the human STARR-seq vector (linearized with *Age*I and *Sal*I) with the NEBuilder HiFi DNA Assembly Cloning Kit (NEB), following the manufacturer’s protocol. We transformed 10-beta electrocompetent cells (NEB C3020) with the plasmids in duplicate following the manufacturer’s protocol, along with a no insert negative control. We pooled the two transformations during recovery and plated 15 µL to estimate complexity. The following day, we estimated complexity as approximately 750,000, and grew a third of the transformation to represent a library of 250,000 in 100 mL of LB+Ampicillin, so that each candidate enhancer is expected to associate on average with 100 different barcodes. We extracted the plasmid using the ZymoPURE II Plasmid Midiprep Kit (Zymo Research).

### Barcode association library for the 9 MPRA assays

We amplified 5 ng of the human STARR-seq (HSS) library with the following primers: P5-STARR-AG-ass-f and P7-STARR-ass-r (**Supplemental Table 8**). These primers add a sample-specific barcode and Illumina flow cell adapters. We then spiked the library into a NextSeq Mid 300 cycle kit with paired-end 149 bp reads and a 20 bp index read (which captured the 15 bp barcode as well as 5 bp of extra sequence to help filter for read quality), using the following custom primers: Read1 as STARR-AG-seq-R1, Read2 as spacer-seq-R2, Index1 as pLSmp-ass-seq-ind1 and Index2 as STARR-AG-ind2 (**Supplemental Table 8**).

### Library cloning

#### From HSS to ORI vector

We amplified 5 ng of the HSS library with the following primers: STARR-Seq-AG-f and STARR-Seq-AG-r (**Supplemental Table 8**) using KAPA HiFi 2x Readymix (Kapa Biosystems) with a 65 °C annealing temperature and 30 s extension. These primers amplify both candidate enhancers and previously assigned degenerate barcodes and add homology arms to the ORI vector (Addgene 99296)^25^. We followed the reaction in real time with Sybr Green (Thermo fisher scientific) and stopped the reaction before plateauing, at 13 cycles. We gel extracted the amplicon on a 1.5% agarose gel as described above. We then cloned the library in a 2:1 molar excess into 100 ng of the hSTARR-seq_ORI vector (addgene ID:99296), linearized with *Age*I and *Sal*I, using the NEBuilder HiFi DNA Assembly Cloning Kit (NEB), following the manufacturer’s protocol. We then transformed 10-beta electrocompetent cells (NEB C3020) with the plasmids in duplicate following the manufacturer’s protocol, along with a no insert negative control. We pooled the two positive transformations during recovery, plated 15 µL to estimate complexity and grew the remainder of the culture in 100 mL LB+Amp. The following day, we estimated the complexity as >500K and extracted the plasmid using the ZymoPURE II Plasmid Midiprep Kit (Zymo Research).

#### From human STARR-seq to pGL4.23c MPRA vector

As described above, we amplified 5 ng of the human STARR-seq library with the following primers: pGL423c-AG-1f and pGL423c-AG-1r (**Supplemental Table 8**). These primers amplify both candidate enhancers and previously assigned degenerate barcodes and add homology arms to the pGL4.23c MPRA vector (GenBank MK484105). We stopped the reaction before plateauing, at 18 cycles. We linearized the pGL4.23c MPRA backbone, while removing the minimal promoter and reporter using *Hin*dIII and *Xba*I. We treated the linearized plasmid with antarctic phosphatase (NEB) following the manufacturer’s protocol, and then gel extracted the plasmid on a 1% agarose gel, as described above. We then cloned our insert into the linearized backbone, transformed, and extracted DNA as described above. We then relinearized the pGL4.23c backbone, containing our enhancer library with *Sbf*I and *Eco*RI, gel extracted, and inserted our minimal promoter + GFP cassette, which contains overlaps for *Sbf*I and *Eco*RI.

#### From human STARR-seq to lentiMPRA 5’/5’

We used similar methods as in the pGL4.23c library cloning with the following changes. The human STARR-seq library was amplified with pLSmP-AG-2f and pLSmP-AG-5r (**Supplemental Table 8**) for 17 cycles. After gel extraction, we cloned the insert into the pLS-mP (Addgene 81225)^13^, which had been linearized with *Sbf*I and *Age*I and treated with antarctic phosphatase. The resulting library was recut with *Sbf*I and *Age*I, residing between the designed candidate enhancer and barcode, and the minimal promoter was ligated in. We generated the minimal promoter with oligos minP_F and minP_R (**Supplemental Table 8**), which provide overlaps for *Sbf*I and *Age*I. The minimal promoter oligos were phosphorylated and annealed using T4 Ligation Buffer and T4 Polynucleotide Kinase (NEB) at 37 °C for 30 minutes followed by 95 °C for five minutes, ramping down to 25 °C at 5 °C/min. We then diluted the annealed oligos at 1:200 and cloned into the linearized pLS-mP backbone with our enhancer library at a 2:1 molar excess.

#### From human STARR-seq to lentiMPRA 5’/3’

We used similar methods as for the pGL4.23c library with the following changes. The human STARR-seq library was amplified with pLSmP-AG-2f and pLSmP-AG-3r (**Supplemental Table 8**) for 17 cycles. After gel extraction, we cloned the insert into the pLS-mP backbone (Addgene 81225)^13^, which had been linearized with *Sbf*I and *Eco*RI, and treated with antarctic phosphatase. Similar to the pGL4.23c library, the resulting library was recut with *Sbf*I and *Eco*RI again, and we inserted our minimal promoter + GFP cassette, containing overlaps for *Sbf*I and *Eco*RI.

#### From human STARR-seq to lentiMPRA 3’/3’

We used similar methods as for he pGL4.23c library with the following changes. The human STARR-seq library was amplified with pLSmP-AG-3f and pLSmP-AG-3R (**Supplemental Table 8**) for 13 cycles. After gel extraction, we cloned the insert into the pLS-mP backbone (Addgene 81225)^13^, which had been linearized with *Eco*RI only and treated with antarctic phosphatase.

### Cell culture, lentivirus packaging, and titration

HEK293T and HepG2 cell culture, lentivirus packaging and titration were performed as previously described with modifications^13^. Briefly, twelve million HEK293T cells were seeded in 15 cm dishes and cultured for 48 hours. To generate wild-type lentiviral libraries (5’/5’ WT, 5’/3’ WT, and 3’/3’ WT), the cells were co-transfected with 5.5 μg of lentiMPRA libraries, 1.85 μg of pMD2.G (Addgene 12259) and 3.65 μg of psPAX2 (Addgene 12260), which encodes a wild type *pol*, using EndoFectin Lenti transfection reagent (GeneCopoeia) according to manufacturer’s instruction. To generate non-integrating lentiviral libraries (5’/5’ MT, 5’/3’ MT, and 3’/3’ MT), pLV-HELP (InvivoGen) that encodes a mutant *pol* was used instead of psPAX2. After 18 hours, cell culture media was refreshed and TiterBoost reagent (GeneCopoeia) was added. The transfected cells were cultured for 2 days and lentivirus were harvested and concentrated using the Lenti-X concentrator (Takara) according to manufacturer’s protocol. To measure DNA titer for the lentiviral libraries, HepG2 cells were plated at 1×10^5^ cells/well in 24-well plates and incubated for 24 hours. Serial volume (0, 4, 8, 16 μL) of the lentivirus was added with 8 μg/ml polybrene, to increase infection efficiency. The infected cells were cultured for three days and then washed with PBS three times. Genomic DNA was extracted using the Wizard SV genomic DNA purification kit (Promega). Multiplicity of infection (MOI) was measured as relative amount of viral DNA (WPRE region, WPRE_F and WPRE_F) over that of genomic DNA [intronic region of *LIPC* gene, LIPC_F and LIPC_R (**Supplemental Table 8**)] by qPCR using SsoFast EvaGreen Supermix (BioRad), according to manufacturer’s protocol.

### Transient transfections and lentiviral infections

HepG2 cells were seeded in 10 cm dishes (2.4 million cells per dish) and incubated for 24 hours. For plasmid-based MPRA, the cells were transfected with 10 μg of the plasmid libraries (HSS, ORI, and pGL4) using X-tremeGENE HP (Roche) according to the manufacturer’s protocol. The X-tremeGENE:DNA ratio was 2:1. For the lentiMPRA, the cells were infected with the lentiviral libraries (5’/5’ WT/MT, 5’/3’ WT/MT, and 3’/3’ WT/MT) along with 8 μg/ml polybrene, with the estimated MOI of 50 for wild-type and 100 for mutant libraries. The cells were incubated for 3 days, washed with PBS three times, and genomic DNA and total RNA was extracted using AllPrep DNA/RNA mini kit (Qiagen). mRNA was purified from the total RNA using Oligotex mRNA mini kit (Qiagen). All experiments for nine libraries were carried out simultaneously to minimize batch effect. Three independent replicate cultures were transfected or infected on different days.

### RT-PCR, amplification, and sequencing of RNA and DNA

DNA for all experiments was quantified using the Qubit dsDNA Broad Range Assay kit (Thermo Fisher Scientific). For all samples, a total of 12 µg of DNA was split into 24 50 µL PCR reactions (each with 500 ng of input DNA) with KAPA2G Robust HostStart ReadyMix (Kapa Biosystems) for three cycles with a 65 °C annealing and 40 s extension, using an indexed P5 primer and a unique molecular identifier (UMI)-containing P7 primer (**Supplemental Table 9**). After three cycles, reactions were pooled and purified with a 1.8x AMPure cleanup, following manufacturer’s instructions, and eluted in a total of 344 µL of Qiagen EB. The entire purified product was then used for a second round of PCR, split into 16 × 50 µL reactions each, with primers P5 and P7 (**Supplemental Table 8**). The reaction was followed in real time with Sybr Green (Thermo Fisher Scientific) and stopped before plateauing. PCRs were then pooled and 100 µL of the pooled PCR products was purified with a 0.9x AMPure cleanup and eluted in 30 µL for sequencing.

mRNA for all experiments was treated with Turbo DNase (Thermo Fisher Scientific) following the manufacturer’s instructions and then quantified using the Qubit RNA Assay kit (Thermo Fisher Scientific). For all samples, we performed three 20 µL reverse transcription reactions, each with one third of the sample (up to 500 ng of mRNA). RT was performed using SuperScript IV (Thermo Fisher Scientific) and a gene-specific primer, which attached a UMI (**Supplemental Table 9**), following manufacturer’s instructions.

cDNA for each sample was split into eight 50 µL PCRs using an indexed P5 primer and P7 (**Supplemental Table 8**) for three cycles. Reactions were then pooled together and purified with a 1.5x AMPure reaction and eluted in 129 µL of Qiagen EB. The purified PCR product was then split into six 50 uL PCRs with P5 and P7 following in real time with Sybr Green and stopped before plateauing. PCRs were then pooled and 100 µL of the pooled PCR products was purified with a 0.9-1.8x AMPure cleanup depending on background banding, and eluted in 30 µL for sequencing.

Two experiments at a time (each with three DNA replicates and three RNA replicates) were run on a 75 cycle NextSeq 550 v2 High-Output kit with custom primers for each assay (**Supplemental Table 8**).

### MPRA to evaluate the impact of enhancer orientation

To test enhancers in both orientations relative to the promoter (in the “forward” and “reverse” orientations), we synthesized the same 2,236 genomic sequences tested above^13^, along with 100 controls previously tested in STARR-seq, which are described below in the Length section below^15^. These sequences were synthesized as 192 bp fragments with HSS-F-ATGC and HSS-R (**Supplemental Table 8**). The forward orientation was amplified in a 50 µL PCR reaction using KAPA HiFi 2x Readymix (Kapa Biosystems) and primers “HSS_pGL4_F” and “HSS_pGL4_R1”; the PCR for the reverse orientation used the primers “HSS_pGL4_F_orr2” and “HSSpGL4_1_orr2” (**Supplemental Table 8**). PCRs were followed in real time with Sybr Green, stopped before plateauing (7 cycles), and purified in a 1X AMPure reaction, eluting in 25 µL of Qiagen EB. 1 µL of the purified products were put into a second PCR reaction, which added 15 bp of barcode sequence and homology to the pGL4.23c vector; the forward orientation used primers HSS_pGL4_F and HSS_pGL4_R2, and the reverse orientation used primers HSS_pGL4_F_orr2 and HSS_pGL4_R2 (**Supplemental Table 8**).

We linearized the pGL4.23c MPRA backbone with *Hin*dIII and *Xba*I (removing the minimal promoter and reporter), and gel extracted the backbone and insert PCR products. Inserts were cloned into the pGL4.23c plasmid using NEBuilder HiFi DNA Assembly Cloning Kit (NEB), following the manufacturer’s protocol. We transformed 10-beta electrocompetent cells (NEB C3020) with the plasmids, grew up transformations in 100 mL of LB+Amp, and extracted plasmid libraries using a ZymoPure II Plasmid Midiprep Kit (Zymo Research).

To clone in the minimal promoter and GFP for the forward orientation, 20 ng of the forward backbone was amplified with Len_lib_linF and Len_lib_linR (**Supplemental Table 8**) using NEBNext High-Fidelity 2X PCR Master Mix (NEB); the minimal promoter and GFP was amplified from 10 ng of the pLS-mP plasmid using minGFP_Len_HAF and minGFP_Len_HAR (**Supplemental Table 8**). For the reverse orientation, 20 ng of the backbone was linearized with Len_lib_linF and Rorr_R2_LinR (**Supplemental Table 8**); for the reverse orientation insert, previously gel extracted minimal promoter and GFP from pLS-mP was amplified using minPGFP_Revorr_Len_HA_F and Len_lib_linR (**Supplemental Table 8**). Both backbones were treated with antarctic phosphatase, following manufacturer’s protocol. All backbones and inserts were gel extracted, with the exception of the reverse orientation insert, which we purified in a 1.8x AMPure reaction. Plasmid libraries were cloned and extracted as previously described.

Transfections (4 independent transfections), DNA/RNA extractions, reverse transcription of mRNA, and qPCRs to amplify barcodes for sequencing were all performed as previously described for the enhancer-length experiments. The final PCRs for the DNA samples were purified in a 1.5X AMPure reaction, using 50 µL of PCR reaction, and eluting in 15 µL of Qiagen EB; cDNA PCRs were gel purified. Libraries were separately denatured and pooled, pooling twice as much of the RNA samples as the DNA samples. Samples were loaded at a final concentration of 1.8 pM on a 75 Cycle NextSeq v2 High-Output kit.

### MPRA to evaluate the impact of including additional sequence context at tested elements

#### Design of enhancer length libraries for array synthesis

We chose to synthesize the same 2,236 genomic sequences tested above^13^. We also included the top 50 and bottom 50 haplotypes, averaging 409 bp, from a screen conducted in the STARR-seq vector^15^ and designed libraries of 192 bp and 354 bp sequences, centered at the position of the previously tested design. We also designed a library of 678 bp sequences for the 2,236 genomic sequences above. We extracted genomic sequence using bedtools getfasta^34^. To the 192 bp library, we added the HSS-F-ATGC sequence to the 5’ end and the HSS-R-clon sequence to the 3’ end (**Supplemental Table 8**).

For the 354 bp library, we split each sequence into two overlapping fragments, A and B. Fragment A included positions 1-190 and fragment B included positions 161-354. To fragment A, we appended the HSS-F-ATGC adapter to the 5’ end and the DO_15R_Adapter to the 3’ end. To fragment B, we appended the DO_5F_Adapter to the 5’ end and the HSS-R-clon adapter to the 3’ end (**Supplemental Table 8**).

For the 678 bp library, we only designed the 2,236 sequences from Inoue *et al.*^13^. We split the sequences into 13 different sets of 172 sequences each. We then split each sequence into four fragments. Fragment A included positions 1-190, fragment B included positions 161-352, fragment C included positions 323-514, and fragment D included positions 485-678. Adapters and primers used for the 13 sets of HMPA are included in **Supplemental Table 10**.

#### Amplification of the 192 bp library

All 192 bp enhancers were amplified from the array using HSSF-ATGC and HSS-R-clon (**Supplemental Table 8**) with KAPA HiFi HotStart Uracil+ ReadyMix PCR Kit (Kapa Biosystems) with SYBR Green (Thermo Fisher Scientific) on a MiniOpticon Real-Time PCR system (Bio-Rad), and stopped before plateauing.

#### Multiplex pairwise assembly for 354 bp library

All 5’ fragments were amplified off the array using HSSF-ATGC and DO_15R_PU (**Supplemental Table 8**) with KAPA HiFi HotStart Uracil+ ReadyMix PCR Kit (Kapa Biosystems) and stopped before plateauing. All 3’ fragments were amplified off the array using DO_5F_PU and HSS-95R (**Supplemental Table 8**). Both were purified using a 1.8x AMPure cleanup and eluted in 20 µL Qiagen EB. 2 µL of USER enzyme (NEB) was added directly to each purified PCR product, and incubated for 15 minutes at 37 °C followed by 15 minutes at room temperature. Reactions were then treated with the NEBNext End Repair Module (NEB) following manufacturer’s protocol, and purified using the DNA Clean and Concentrator 5 (Zymo Research) and eluted in 12 µL EB, following manufacturer’s protocol. We then quantified DNA concentrations for both treated samples using a Qubit and diluted samples to 0.75 ng/uL. We then assembled the 5’ and 3’ fragments as described previously^23^. Briefly, fragments were allowed to anneal and extend for 5 cycles with KAPA HiFi 2X HotStart Readymix (Kapa Biosystems) before primers HSSF-ATGC and DO_95R were added for amplification (**Supplemental Table 8**).

#### Hierarchical multiplex pairwise assembly for 678 bp library

All libraries were amplified off the array using the primers indicated in **Supplemental Table 10** with KAPA HiFi HotStart Uracil+ ReadyMix PCR Kit (Kapa Biosystems) as described above. During the first round of assembly, fragments A and B were assembled with HSSF-ATGC and DO_31R_PU and fragments C and D were assembled with DO_8F_PU and HSS_R (**Supplemental Table 10**). Assembled libraries were then purified with a 0.65x Ampure cleanup following the manufacturer’s protocol, and eluted in 20 µl. 2 µl of USER enzyme (NEB) was added to the purified assembly reactions and incubated at 37 °C for 15 minutes followed by 15 minutes at room temperature, and then repaired using the NEBNext End Repair Module (NEB), following manufacturer’s protocol, and purified using the DNA Clean and Concentrator 5 (Zymo Research) and eluted in 10 µL EB. All libraries were then quantified using the Qubit dsDNA HS Assay kit (Thermo Fisher Scientific) and eluted to 0.75 ng/ul. Assemblies AB and CD were then assembled together following the multiplex pairwise assembly protocol^23^. After the second assembly, libraries were purified using a 0.6x AMPure cleanup and eluted in 30 µL EB. We then amplified 1 uL of each assembly with HSS-F-ATGC-pu1F and HSS-R-clon-pu1R to add flow cell adapters and indexes (**Supplemental Table 8**). We performed the assembly for each set of 172 sequences separately, as well as for different combinations of sets, up to all 2,236 sequences at once.

#### Sequence validation of assembled libraries

Before cloning, we verified assembly and uniformity of our libraries. The multiplex pairwise assembly library (2,336 354mers) was sequenced on a Miseq v3 600 cycle kit with paired-end 305 bp reads. Reads were merged with PEAR v0.9.5^35^ and aligned to a reference fasta file with BWA mem^36^. Each of the 13 hierarchical pairwise assembly sub-libraries (172 678mers) as well as different complexities (344, 688, 1032, 1376, 1720, 2064, 2236) were sequenced on a Miseq v3 600 cycle kit with paired-end 300 bp reads. Paired end reads were aligned to a reference fasta file with BWA mem^36^. As our HMPA library was longer than the maximum Illumina sequencing length (600 bp), we prepared our HMPA sub-library 3 (172 678mers) for sequencing on the PacBioSequel System using V2.1 chemistry (Pacific Biosciences). The library was amplified with pu1L and pu1R and sent to the University of Washington PacBio Sequencing Services for library preparation and sequencing. We obtained 312,277 productive ZMWs with an average Pol Read length of 30,806 bp. After generating circular consensus sequences, we obtained 218,240 CCS reads with a mean read length of 882 bp.

#### Barcoding and cloning of length libraries into pGL4.23c

We performed a two-step PCR to add barcodes and cloning adapters for pGL4.23c onto our three different libraries. For the 192mer and 354mer library, we amplified 20 ng of the library with HSS-pGL4_F and HSS-pGL4_R1 (**Supplemental Table 8**) using NEBNext High-Fidelity 2X PCR Master Mix (NEB) for 16 cycles. For the 678mer libraries, we pooled all 13 sub-libraries at equal concentrations, and then amplified 20 ng with the same primers above and conditions above. All PCR products were purified with a 1.5x AMPure cleanup following manufacturer’s instructions and eluted in 50 µL. We then used 1µL of each purified reaction for a second PCR to append the 15 bp degenerate barcodes and cloning adapters. For the second reaction, we used HSS-pGL4_F and HSS_pGL4_R2 (**Supplemental Table 8**).

We linearized the pGL4.23c MPRA backbone, while removing the minimal promoter and reporter using *Hin*dIII and *Xba*I. We treated the linearized plasmid with antarctic phosphatase following the manufacturer’s protocol, and then gel extracted the plasmid on a 1% agarose gel. We then cloned all three libraries into the pGL4.23c plasmid using the NEBuilder HiFi DNA Assembly Cloning Kit (NEB), following the manufacturer’s protocol. The library was then transformed into 10-beta electrocompetent cells (NEB C3020), grown in 100 mL of LB+Amp, and extracted using the ZymoPure II Plasmid Midiprep Kit (Zymo Research). We then relinearized each library with Len_lib_linF and Len_lib_linR and amplified the minimal promoter and GFP from 10 ng of the pLSMP plasmid using minGFP_Len_HAF and minGFP_Len_HAR (**Supplemental Table 8**). We then gel extracted all linearized libraries and the minimal promoter + GFP insert on a 1% agarose gel. We inserted the minimal promoter and GFP using the NEBuilder HiFi DNA Assembly Cloning Kit (NEB) as described above.

#### MPRA of all enhancer length libraries

The day before transfection, we seeded HepG2 cells in five 10 cm dishes. Day of transfection, we combined the 192, 354, and 678 pGL4.23c libraries at a 1:1:1 molar ratio and transfected 21 µg of pooled libraries into each 10 cm dish using Lipofectamine 3000 (Thermo Fisher Scientific), following the manufacturer’s protocol. 48 hours post transfection, we extracted DNA and RNA from each replicate using the AllPrep DNA/RNA Mini Kit (Qiagen), following manufacturer’s instructions.

We added UMIs to a total of 4 µg of DNA from each replicate split across eight reactions with KAPA2G Robust HotStart ReadyMix (Kapa Biosystems) for three cycles with a 65 °C annealing and 40 s extension, using P5-pLSmP-5bc-idx and P7-pGL4.23c-UMI (**Supplemental Table 8**). After three cycles, reactions were pooled and purified with a 1.8x AMPure cleanup, following manufacturer’s instructions, and eluted in a total of 87 µL of Qiagen EB. The entire purified product was then used for a second round of PCR, split into 6 50 µL reactions each, with primers P5 and P7. The reaction was followed in real time with Sybr Green and stopped before plateauing. PCRs were then pooled and 100 µL of the pooled PCR products was purified with a 0.9x AMPure cleanup and eluted in 30 µL for sequencing.

RNA for each replicate was treated with Turbo DNase (Thermo Fisher Scientific) following manufacturer’s protocol and then quantified using the Qubit RNA Assay kit (Thermo Fisher Scientific). For all samples, we performed two 15 µL reverse transcription reactions, using a total of 15.75 µL RNA (½ total). RT was performed using Thermo Fisher SuperScript IV (Thermo Fisher Scientific) and a gene-specific primer (P7-pGL4.23c-UMI), which attached a UMI, following the manufacturer’s instructions. cDNA for each sample was split into four 50 µL PCRs using P5-pLSmP-5bc-idx and P7 for three cycles. Reactions were then pooled together and purified with a 1.5x AMPure reaction and eluted in 64.5 µL of Qiagen EB. The purified PCR product was then split into three 50 µL PCRs with P5 and P7, followed in real time with Sybr Green (Thermo Fisher Scientific), and stopped before plateauing (11 cycles). PCR products were purified with a 1.5x AMPure reaction before sequencing on a 75 cycle NextSeq 550 v2 High-Output kit.

For barcode associations, we amplified 5 ng of each library with P5_pGL4_Idx_assF and P7-pGL4-ass-R (**Supplemental Table 8**), following in real time with Sybr Green for 14-15 cycles. PCR products were purified with a 1x AMPure cleanup and eluted in 20 µL of Qiagen EB for sequencing. Libraries were separately denatured and pooled to account for part of the clustering bias on the NextSeq. We brought the 192 library to a final concentration of 1.65 pM, the 354 library to a final concentration of 2.15 pM, and the 678 library to a final concencentration of 2.9 pM. We then pooled an equal volume of each library and loaded on a 300 cycle NextSeq 550 v2 Mid-Output kit with an 80 bp read 1 and 213 bp read 2 (in order to sequence part of contributing oligos A, C, and D).

### MPRA processing pipeline

#### Reproducible MPRA analysis pipeline implementation

We will provide a suite of tools to fully reproduce the processing of raw MPRA data from this study at the following Github page: https://github.com/vagarwal87/MPRA-pipeline. The pipeline is built on the WDL pipeline scripting language (https://software.broadinstitute.org/wdl/) and utilizes Cromwell, a workflow management system. Cromwell helps to run the pipeline on a variety of platforms, including the Amazon Web Services and Google Cloud computing platforms, or a local computing cluster environment that supports Docker or Singularity. The sections below document the various components of the pipeline, which borrow heavily from our earlier work^13^.

#### Associating barcodes to designed elements

For each of the barcode association libraries, we generated Fastq files with bcl2fastq v2.18 (Illumina Inc.), splitting the sequencing data into an index file delineating the barcode and two paired-end read files delineating the corresponding element linked to the barcode. If the paired-end reads overlapped in sequence, they were merged into one and aligned using BWA mem^36^ to a reference Fasta file comprised of the designed elements (**Supplemental Table 2**). We carried forward the subset of merged reads whose mapped length corresponded to the expected length of the designed element ± 5 bp (i.e., 171 ± 5, 192 ± 5, 354 ± 5, and 678 ± 5, depending on the element size), allowing indels or mismatches. To minimize the impact of sequencing errors, we associated a barcode to an element if: i) the barcode:element pair was sequenced at least three independent times, and ii) ≥90% of the barcode mapped to a single element. These barcode associations were then used as a dictionary to match barcodes detected in the RNA and DNA sequencing libraries in different MPRA designs.

#### Replicates, normalization, and RNA/DNA activity scores

Barcodes were counted for RNA and DNA samples for each MPRA experiment, using UMIs to collapse barcodes derived from the same molecule during PCR, and mapped to the element they were linked to, as identified by the dictionary of barcode:element associations. To normalize RNA and DNA for different sequencing depths in each sample, we followed a nearly identical scheme as one we had previously devised^13^. Briefly, for each replicate of each MPRA design, we first considered the subset of barcodes that were observed for both the RNA and DNA samples of the replicate. We then summed up the counts of all barcodes contributing to each element and computed the normalized counts as the counts per million (cpm) sequenced reads of that library. Finally, we computed enhancer activity scores as log_2_(RNA [cpm]/DNA [cpm]). To account for the differential scale among replicates of each experiment, we divided the RNA/DNA ratios by the median across the replicate value before averaging them. Due to low counts in the initial round of sequencing and poor sample quality, the three replicates from the 5’/3’ MT and 3’/3’ MT were resequenced, and the data from each pair of technical replicates was pooled together across the two independent sequencing runs. Even after pooling, the first replicates of these two assays exhibited poorer inter-replicate concordance than the other replicates (**Figure 2A**, **Supplemental Figure 3**), and thus were excluded during replicate averaging (**Supplemental Table 2**). In practice, this decision very modestly altered the numerical results, and did not change the study’s conclusions.

### Modeling and Analyses

#### Features considered

For each candidate enhancer, we computed a total of 915 features derived from either: i) the sequence itself, or ii) experimentally measured information, computed as a mean signal extracted from the corresponding region of the human genome (**Supplemental Table 3**, with a full list of features and data sources: **Supplemental Table 4**). The sequence-based features represent the conservation of the sequence, general G/C content, predicted chromatin state, and likelihood of binding to an assortment of transcription factors and RNA binding proteins. In contrast, the experimentally derived features represent empirical measurements of chromatin/epigenetic state, binding to transcription factors, or transcriptional activity. The features were derived from custom Perl scripts, the UCSC genome browser^37^, DeepSEA^38^, DeepBind^39^, with epigenomic data derived from the Epigenomics Roadmap Consortium^40^, CAGE data from the FANTOM Consortium^30^, and ChIP-seq data from the ENCODE Consortium^7^.

#### Feature pre-processing

Right-skewed data such as ChIP-seq and CAGE signal were log-transformed to approximate a normal distribution, and each feature was then z-score normalized to scale the features similarly. This enabled a direct comparison of coefficients among features derived from the resulting linear models.

#### Model training

As described before^13^, we trained a lasso regression model on each of 10 folds of the data, selecting enhancers which were measured with at least 10 independent barcodes to reduce the impact of measurement noise in the assessment of model quality. A lasso regression model was chosen specifically because it employs an L1 regularization penalty, which leads to the selection of the fewest features that maximally explain the data. The strength of the regularization was controlled by a single *λ* parameter, which was optimized using 10-fold cross-validation on the entire dataset. To evaluate the most relevant features selected, we trained a lasso regression model on the full dataset and visualize the top 30 coefficients with the greatest magnitude. A full table of the selected features and their coefficients are provided (**Supplemental Table 5**).

## SUPPLEMENTAL TABLES

**Supplemental Table 1.** Genomic coordinates (human genome build hg19) and sequences for all designed elements, both naturally occurring as well as synthetic positive and negative controls, in the experiments testing the nine assays, element orientation, and element size.

**Supplemental Table 2.** Activity scores computed for each element for each of the 9 MPRA assays tested. Provided are averaged activity scores across replicates as well as individual scores for each replicate alongside normalized DNA counts, normalized RNA counts, and the number of barcodes per element.

**Supplemental Table 3.**
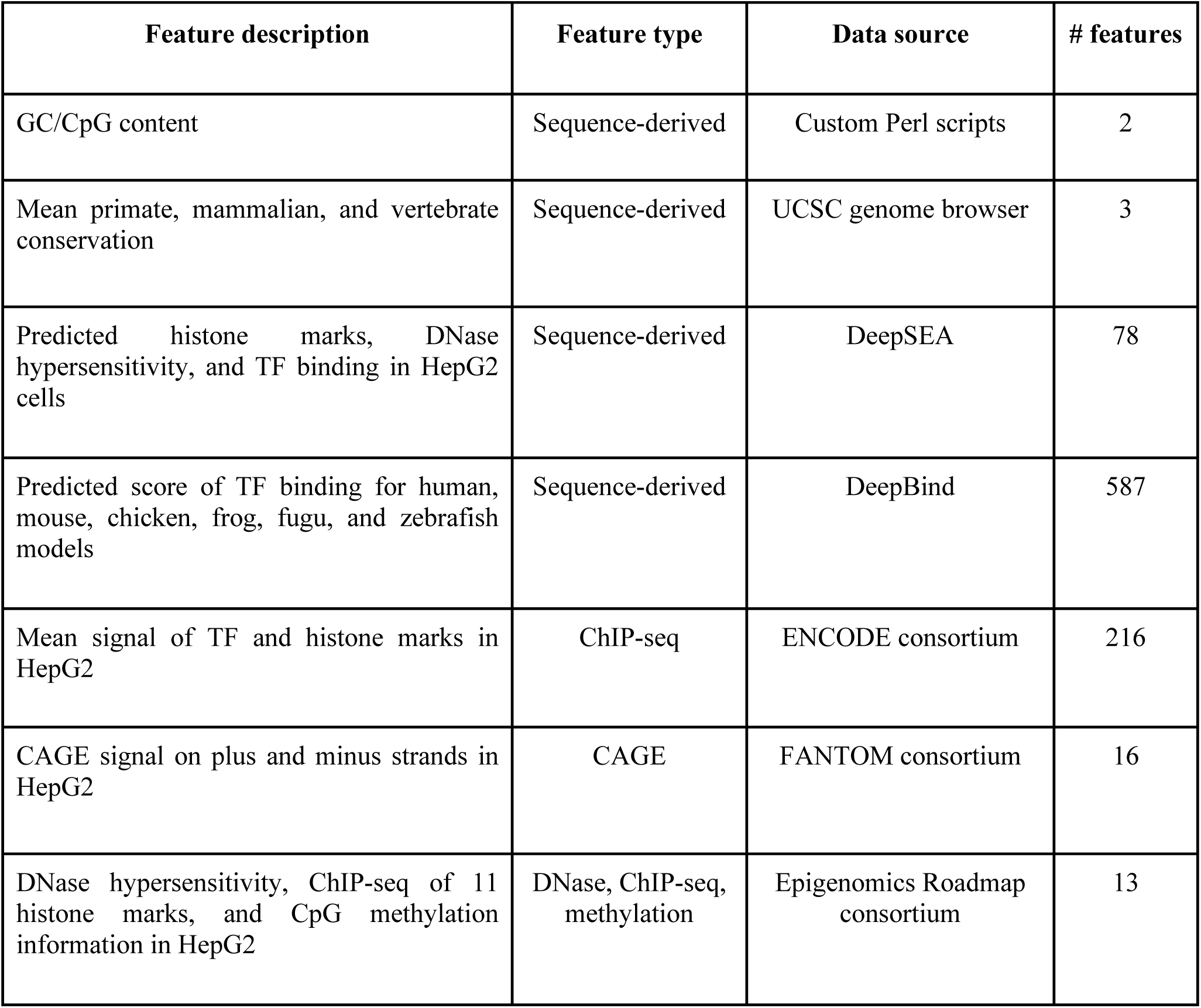
Summary of 915 features considered in a model trained to predict enhancer activity, with an overview of features considered, feature type (i.e., computationally predicted or experimentally derived), data source, and number of features in the category.

**Supplemental Table 4.** Definition of each feature considered in the lasso regression models, with detailed metadata corresponding to the data source of origin, species of origin, sample accession IDs, and additional factor-specific information. Also provided are pre-computed tables of the features used during training for the 9 MPRA assays as well as the assay testing different size classes.

**Supplemental Table 5.** Coefficients fit for the full lasso regression models for each of the 9 MPRA assays, the assay testing different size classes, and differential pairwise comparisons of assays tested therein.

**Supplemental Table 6.** Activity scores computed for each element in the forward (‘F’) and reverse (‘R’) orientations in the orientation assay. Provided are averaged activity scores across replicates as well as individual scores for each replicate alongside normalized DNA counts, normalized RNA counts, and the number of barcodes per element.

**Supplemental Table 7.** Activity scores computed for each element in the short, medium, and long elements in the assay testing for different size classes. Provided are averaged activity scores across replicates as well as individual scores for each replicate alongside normalized DNA counts, normalized RNA counts, and the number of barcodes per element.

**Supplemental Table 8:** All primer, adapter, and oligo sequences utilized throughout the manuscript (excluding HMPA). When applicable, includes the assay and step for which the primer was used.

**Supplemental Table 9.** All sequence indexes used for each experiment.

**Supplemental Table 10.** All primer and adapter sequences used for Hierarchical Multiplex Pairwise Assembly.

## SUPPLEMENTAL FIGURES

**Supplemental Figure 1.**
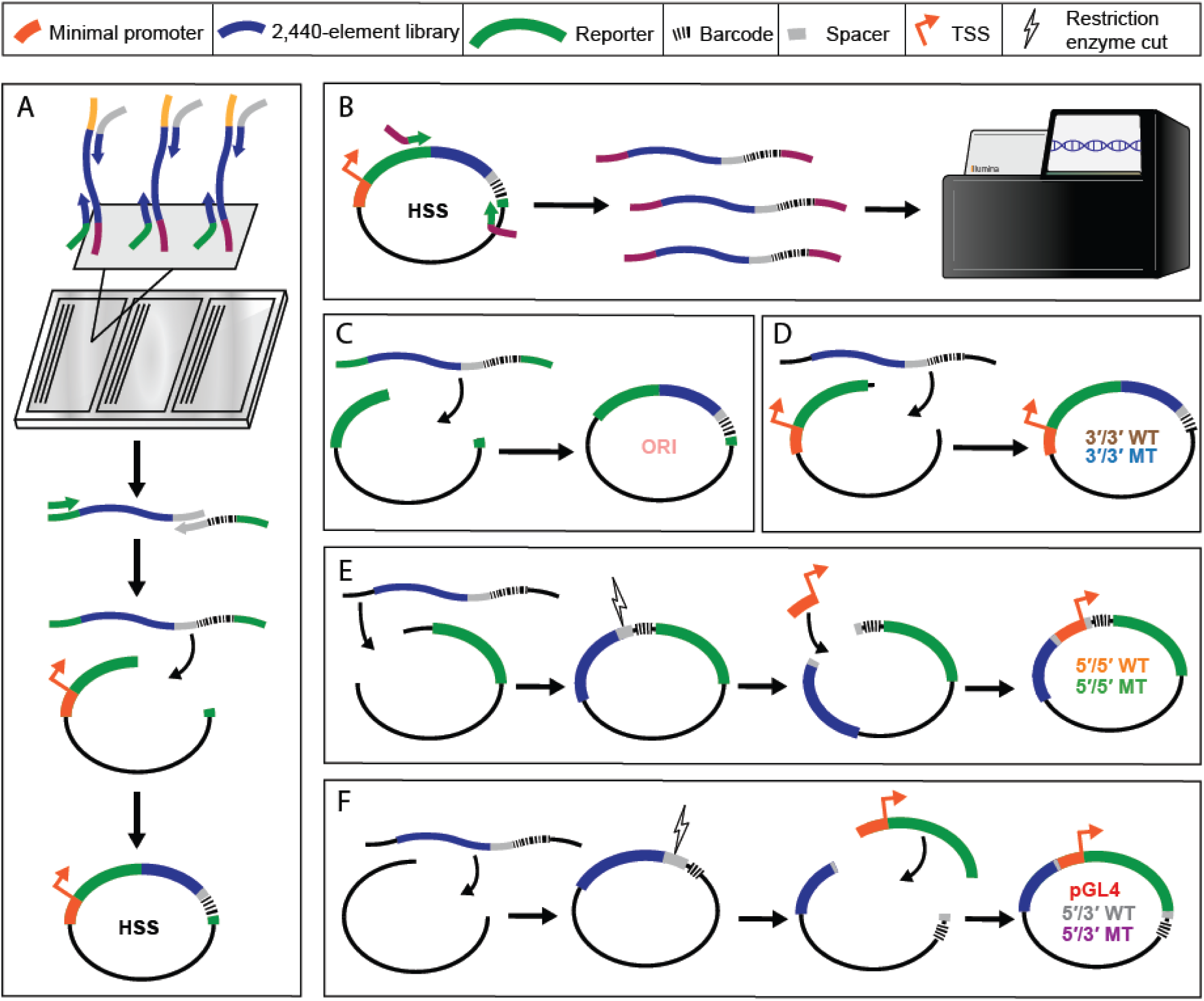
Cloning of MPRA vector libraries. **A)** We amplified an existing array library from Inoue *et al.*^13^, adding homology to the STARR-seq vector on the 5’ end and a spacer sequence for cloning on the 3’ end. The library was put into a second PCR reaction adding barcodes and homology to STARR-seq to the 3’ end, and cloned sequences into the STARR-seq vector. **B)** We amplified enhancers from the STARR-seq vector library, adding flow cell adapters to the sequences, and sequenced the library on a NextSeq Mid 300 cycle kit to associate enhancer sequences to their corresponding barcodes. **C)** The STARR-seq ORI assay was cloned by amplifying enhancers from the STARR-seq library while adding homology to the STARR-seq ORI vector, and cloned into the linearized ORI backbone. **D)** We cloned the 3’/3’ lenti- and mutant lentiMPRA libraries by first linearizing the pLSmP backbone without removing the minimal promoter and GFP. The enhancer library and barcodes were then amplified from the STARR-seq vector adding homology to the linearized backbone, and cloned into the vector. **E)** To clone the 5’/5’ lenti- and mutant lentiMPRA libraries, we linearized the pLSmP vector such that the minimal promoter was removed. We then linearized plasmids at the spacer sequence and cloned in the minimal promoter such that it was between the enhancer and barcode sequences. **F)** The pGL4.23c episomal assay and the 5’/3’ lenti- and mutant lentiMPRA libraries were cloned in the same manner as the 5’/5’ vector library, except the pGL4.23c and pLSmP backbones were initially linearized to remove both the minimal promoter and reporter gene; after cloning in our library, the minimal promoter and reporter were inserted into the spacer sequence such that enhancers were upstream and barcodes downstream of the reporter.

**Supplemental Figure 2.**
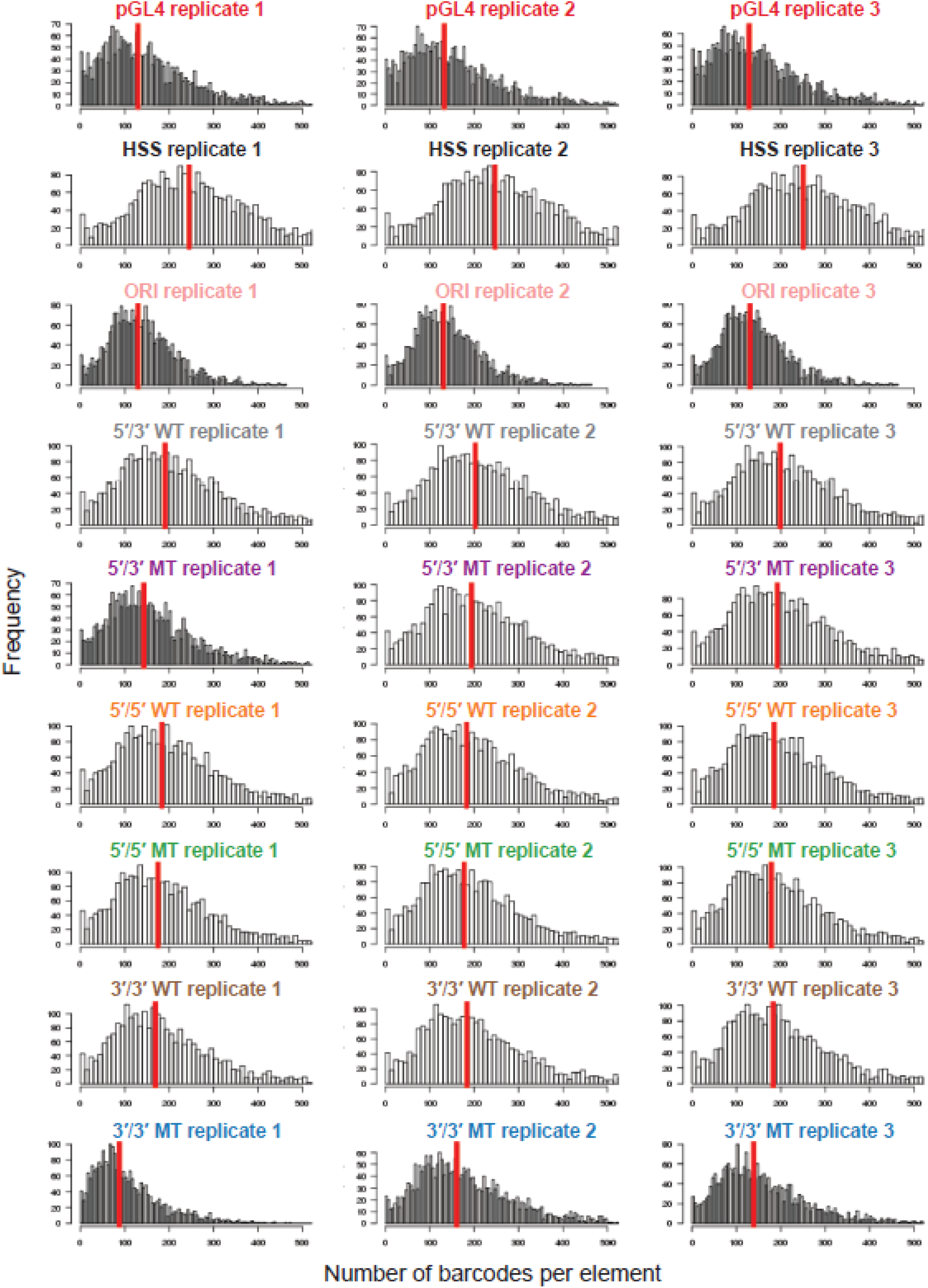
Histograms of barcodes per element for each replicate of the 9 MPRA methods. Shown are histograms indicating the number of observed barcodes per element, for each of the 2,440 elements tested in each of the 3 replicates for each of the 9 MPRA methods. Shown with a vertical red line is the median number of barcodes per element.

**Supplemental Figure 3.**
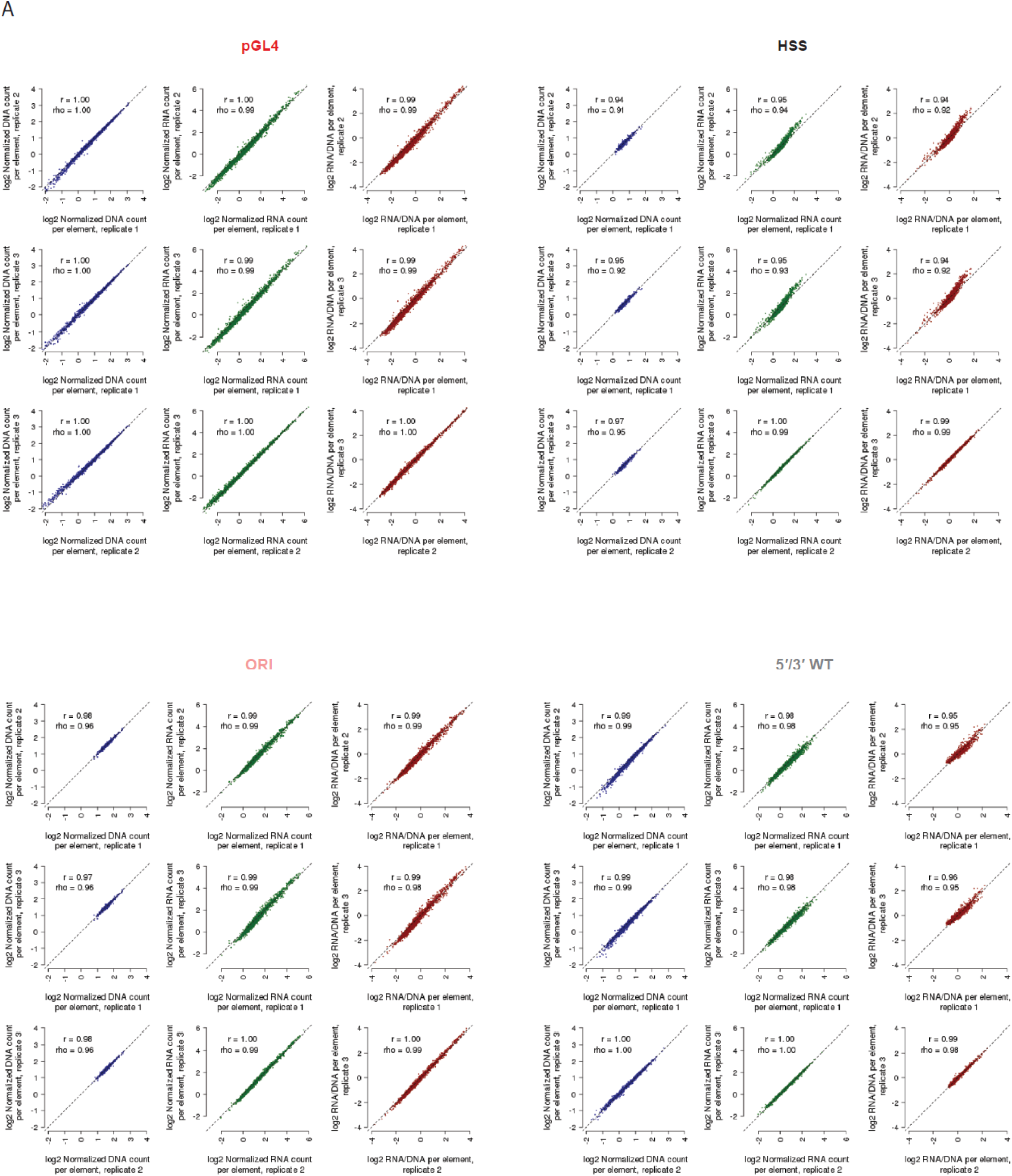
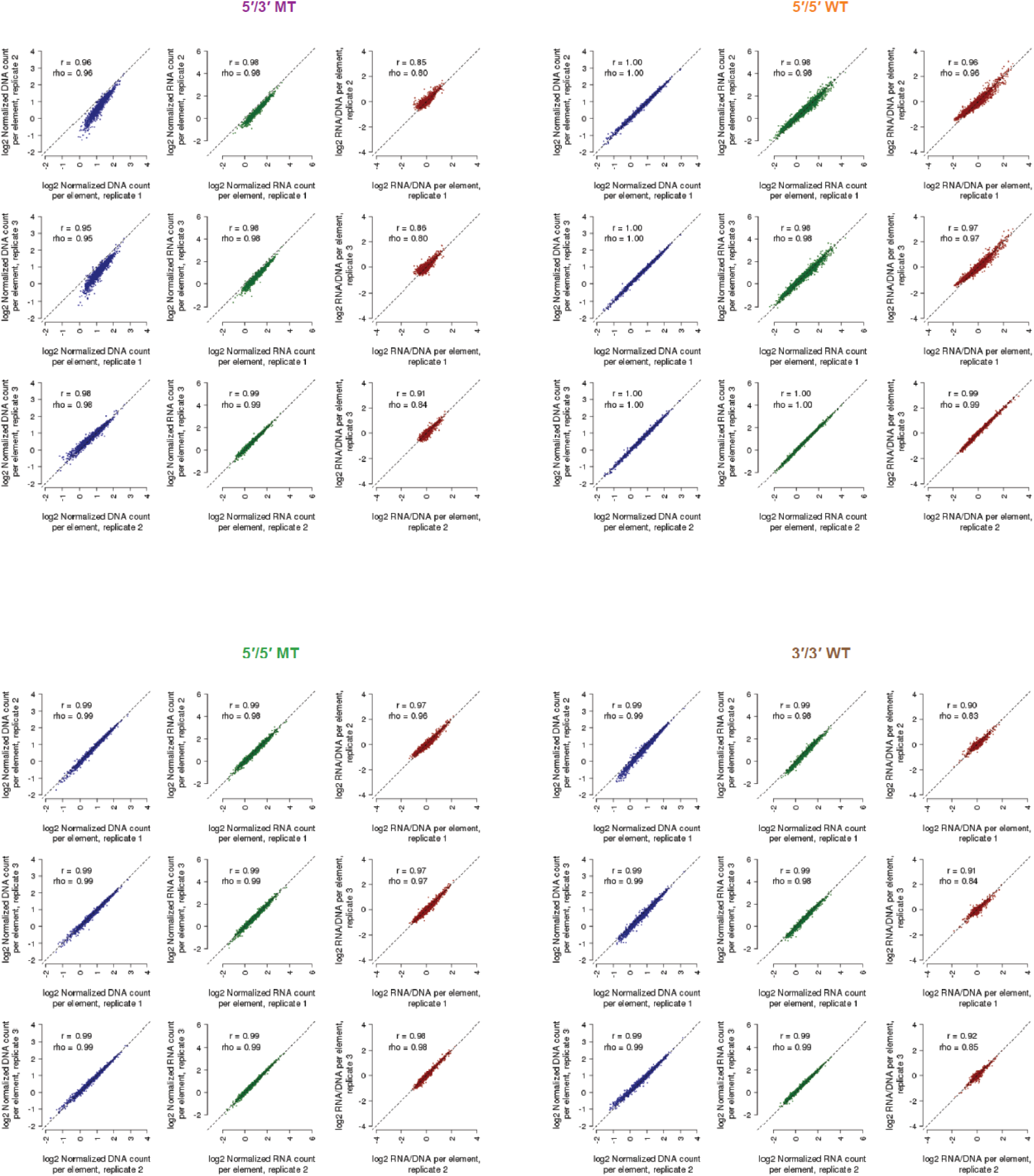
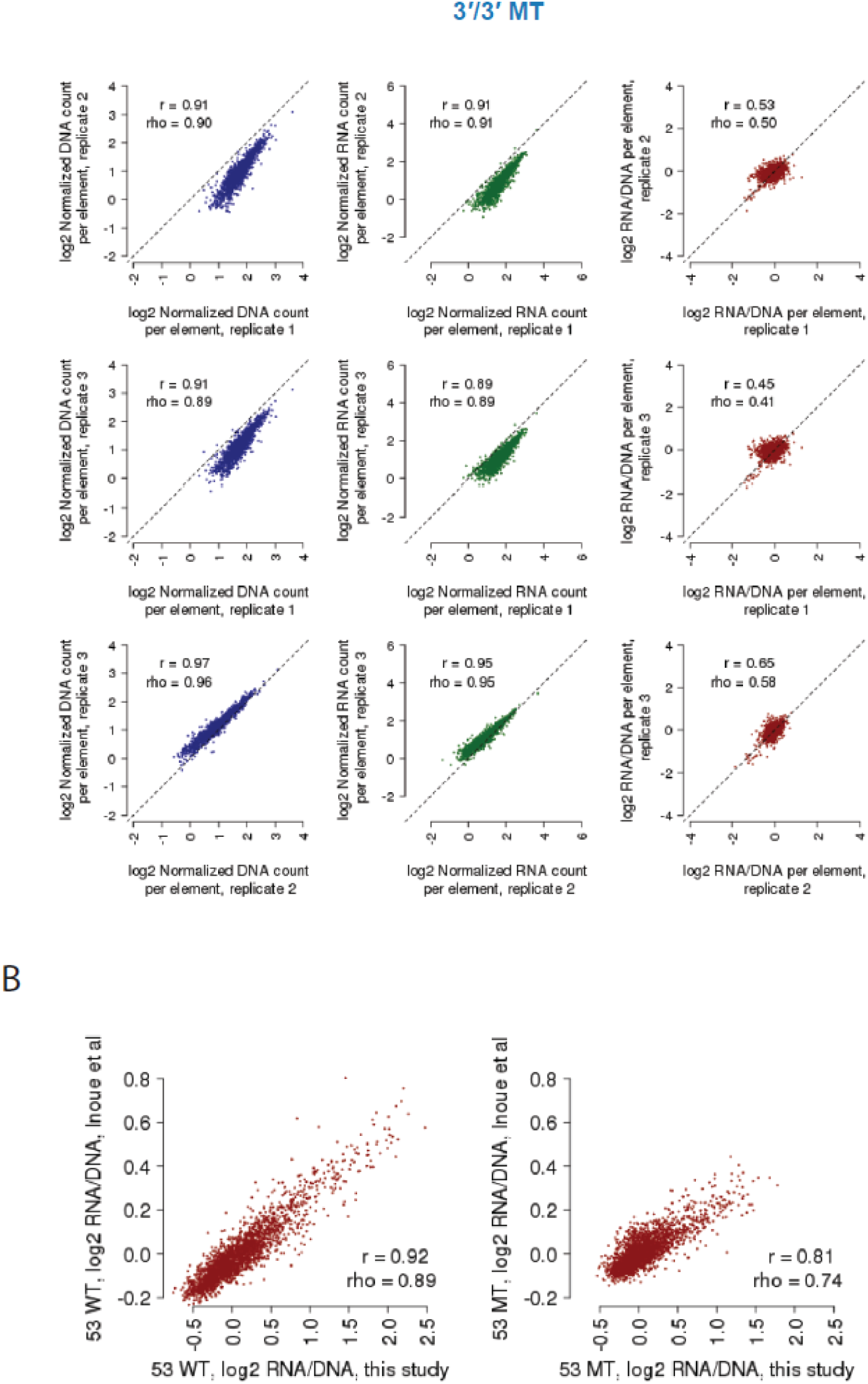
Reproducibility among replicates for each of the 9 MPRA methods. **A)** Shown are scatter plots displaying the relationship between observed DNA counts (blue), RNA counts (green), and RNA/DNA ratios (red) for all pairwise comparisons among replicates, for each of the 9 MPRA methods tested. Also indicated is the Pearson (*r*) and Spearman (rho) correlation values. Candidate enhancers supported by fewer than 10 barcodes were filtered out prior to this analysis to reduce the impact of technical noise. **B)** Scatter plots displaying the relationship between 5’/3’ WT and 5’/3’ MT ratios from this study and the indicated datasets from Inoue *et al*.^13^. Also indicated are the Pearson (*r*) and Spearman (rho) correlation values.

**Supplemental Figure 4.**
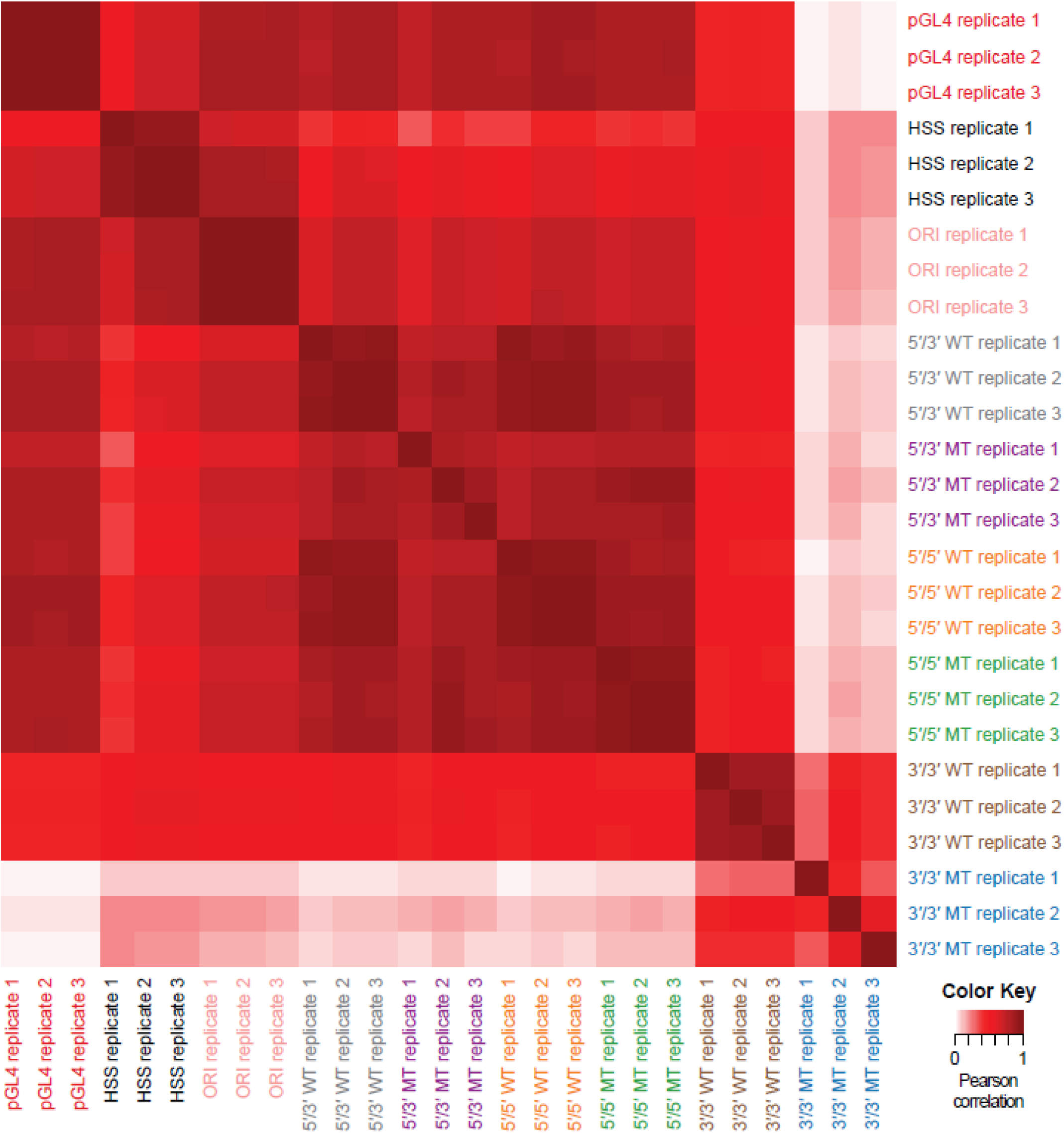
Reproducibility among RNA/DNA ratios among individual replicates for each of the 9 MPRA methods. Shown is a heatmap displaying the pairwise Pearson correlation among individual replicates, using RNA/DNA ratios from each of the 3 replicates for each of the 9 MPRA methods tested.

**Supplemental Figure 5.**
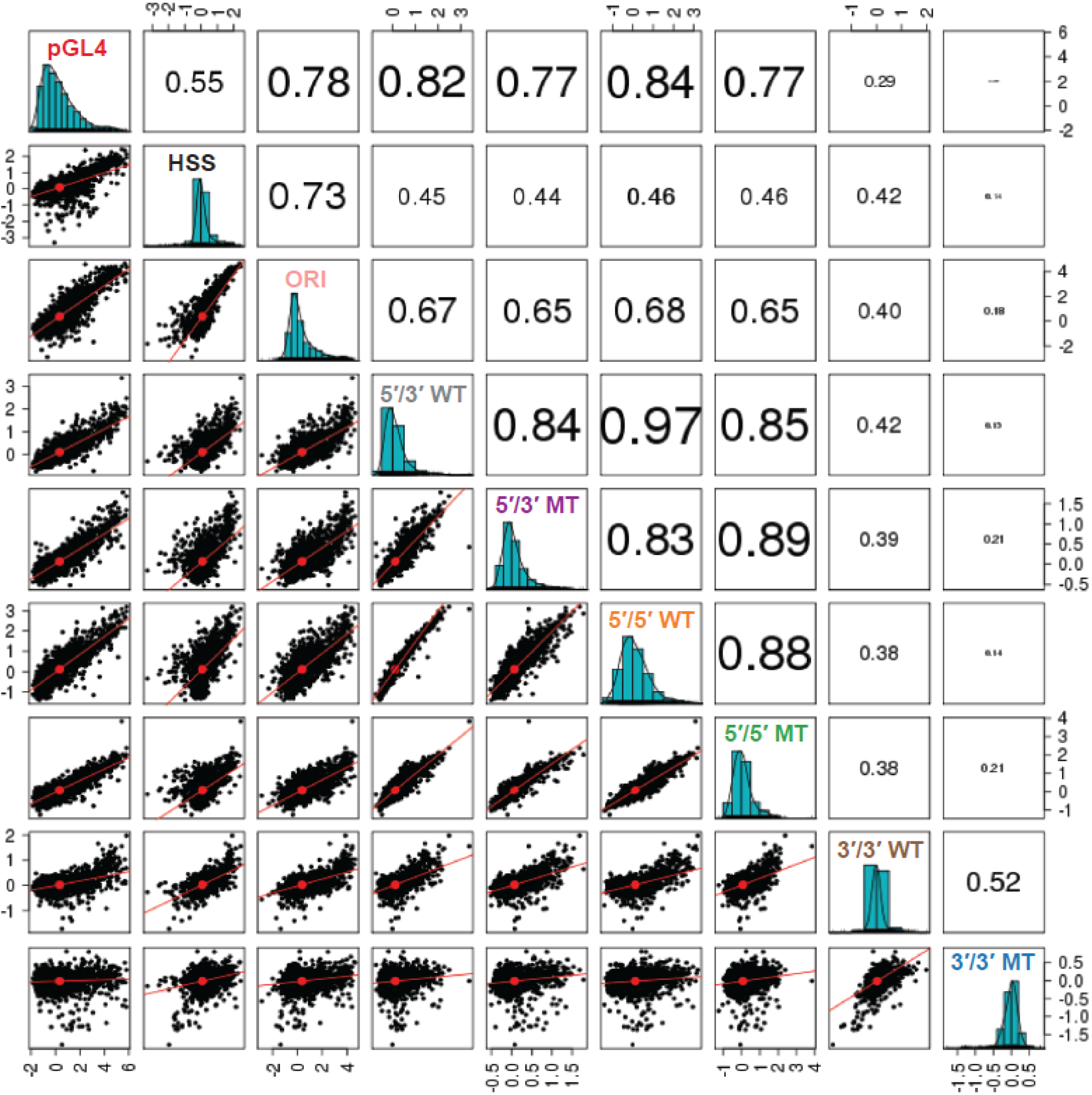
Reproducibility of averaged RNA/DNA ratios. This figure is equivalent to Figure 2B, except it displays the scatter matrix values using Spearman correlation values instead of Pearson correlation values.

**Supplemental Figure 6.**
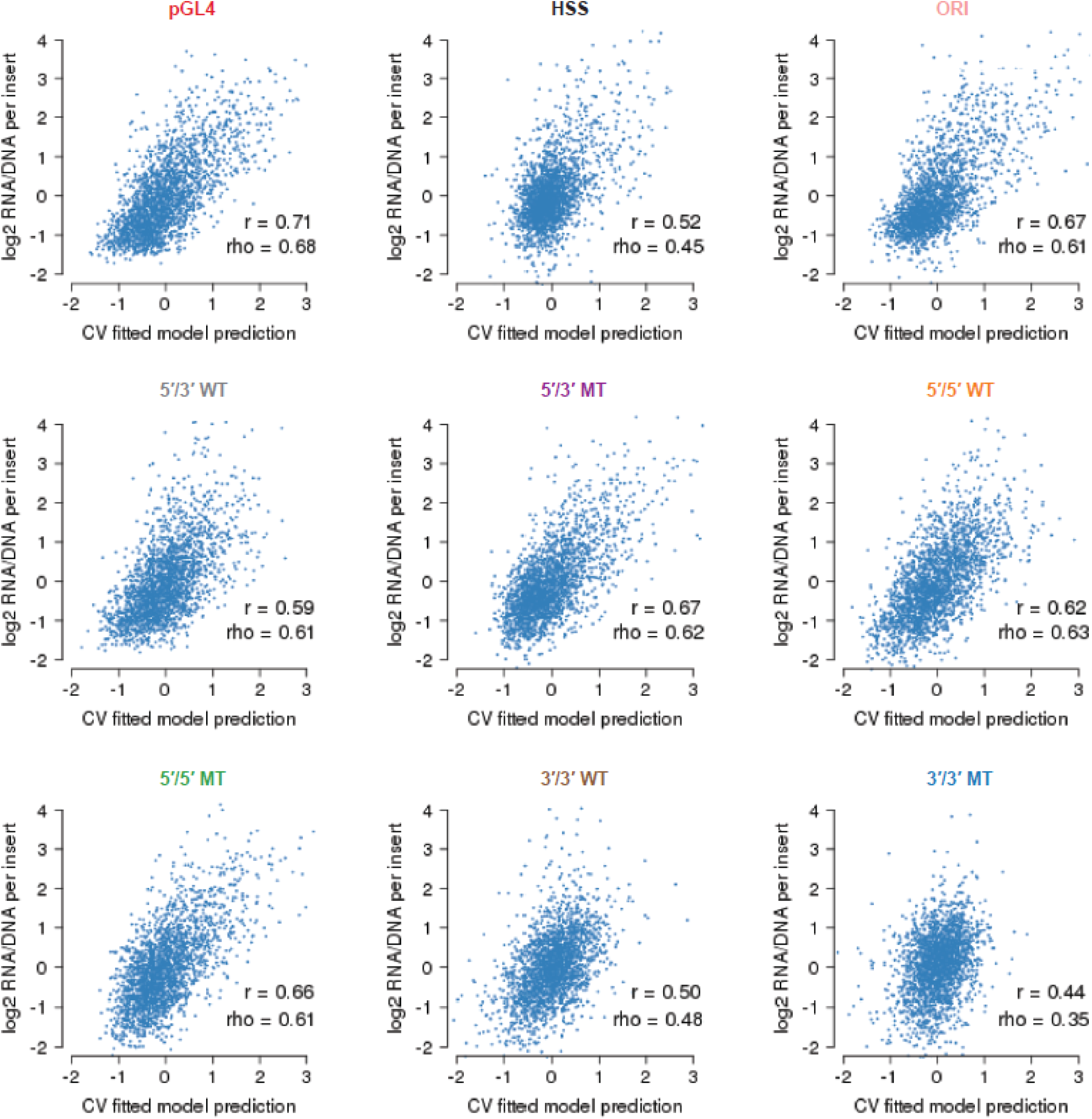
Performance of the predictions on the 9 MPRA methods. Scatter plots displaying the relationship between the 10-fold cross-validated predictions derived from lasso regression models and the observed RNA/DNA ratios, for each of the 9 MPRA methods tested. Also indicated are the Pearson (*r*) and Spearman (rho) correlation values.

**Supplemental Figure 7.**
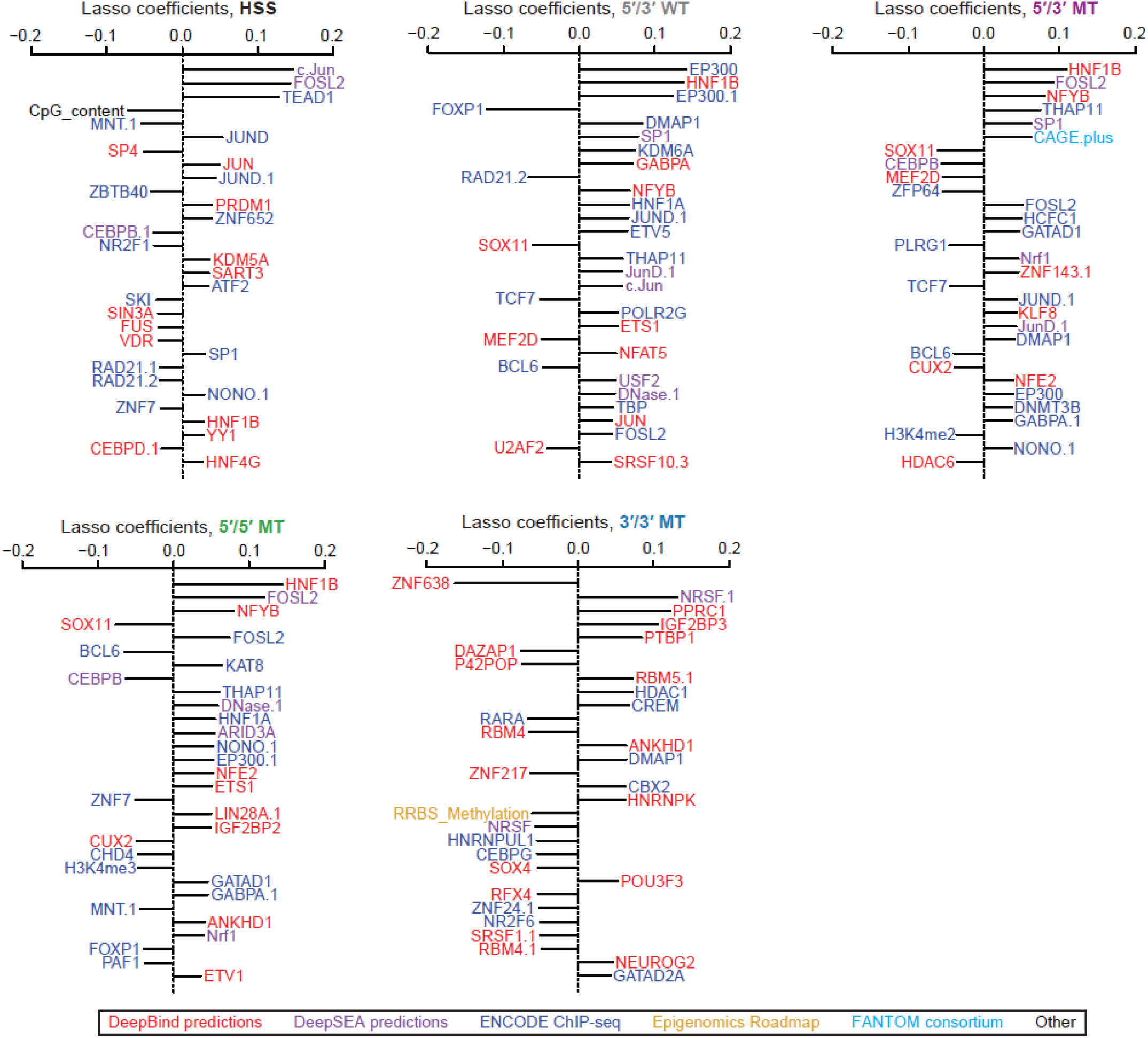
Predictive modeling of the ratios for 5 additional MPRA methods. The top 30 coefficients derived from a lasso regression model trained on the full dataset derived from the five indicated MPRA methods. Features with the extension “.1”, “.2”, etc allude to redundant features or replicate samples. See also **Figure 3B** for the coefficients of the remaining four methods.

**Supplemental Figure 8.**
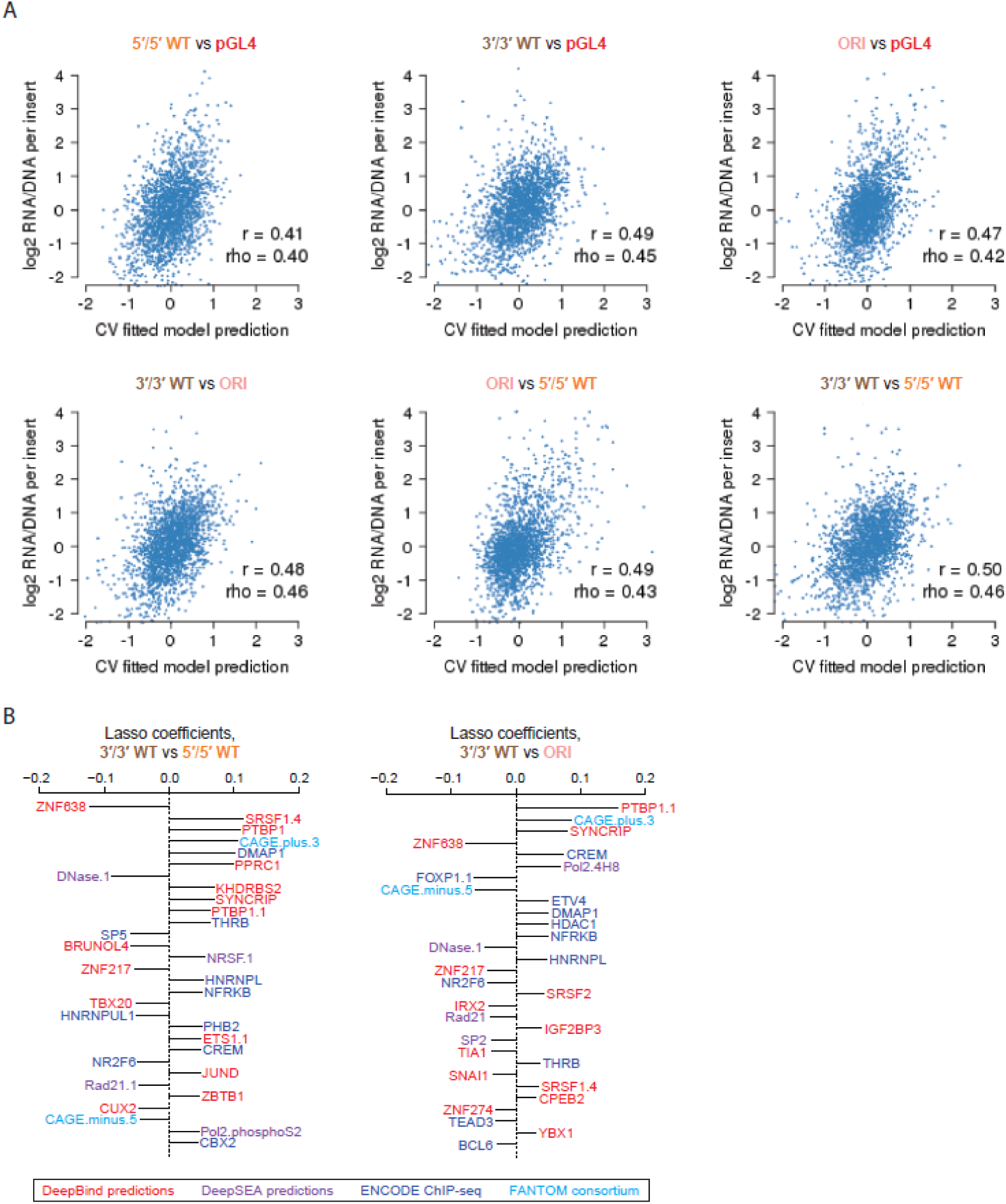
Coefficients and performance of the models to predict differential element efficacy among the 9 MPRA methods. **A)** Scatter plots displaying the relationship between the 10-fold cross-validated predictions derived from lasso regression models and the observed RNA/DNA ratios, for each of the 6 indicated differential comparisons tested. Also indicated are the Pearson (*r*) and Spearman (rho) correlation values. **B)** The top 30 coefficients derived from lasso regression models trained on the full dataset to predict observed differences in the indicated pairs of MPRA methods. Features with the extension “.1”, “.2”, etc allude to redundant features or replicate samples. See also **Figure 3D** for the remaining four comparisons.

**Supplemental Figure 9.**
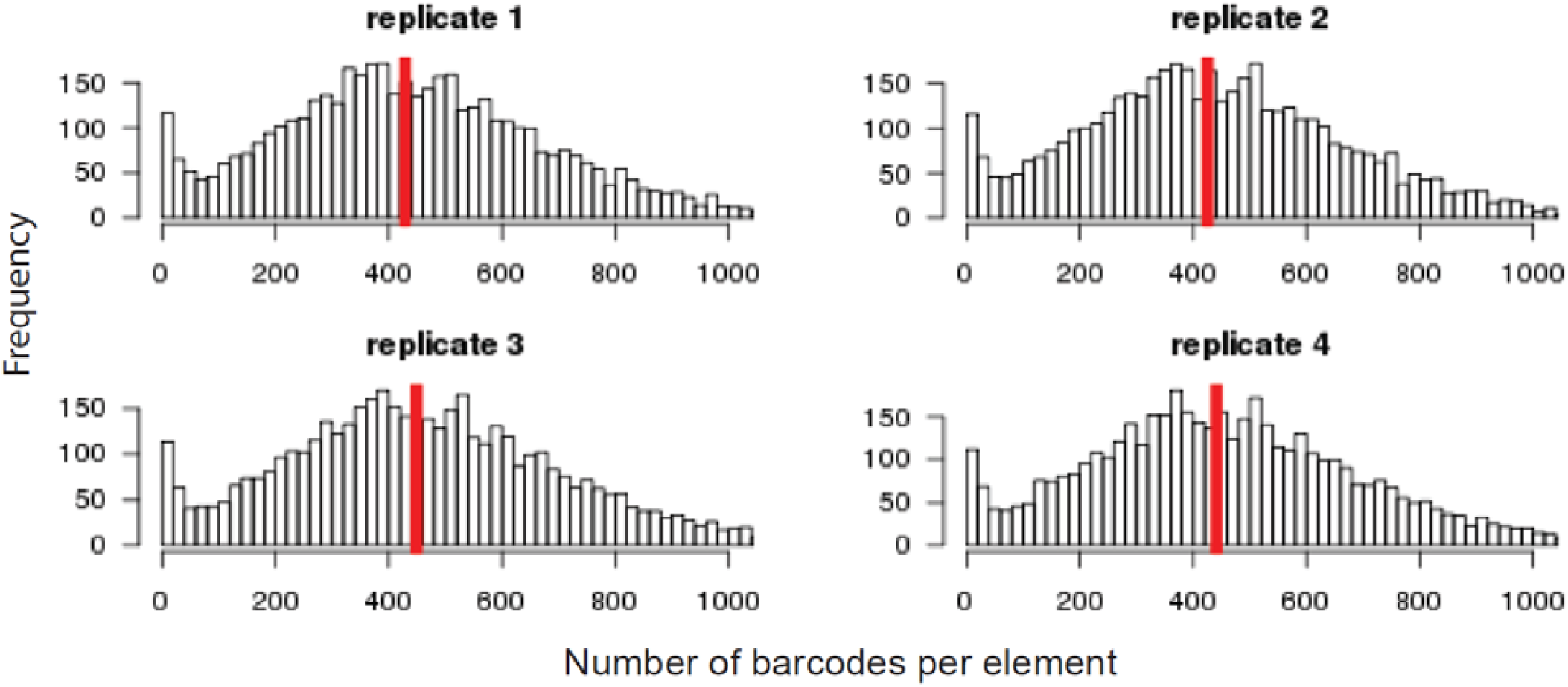
Histograms of barcodes per element for each replicate of the the library testing orientation. Shown are histograms indicating the number of observed barcodes per element, for each of the 2,336 elements tested in each of the 4 replicates for the library testing orientation (i.e., forward and reverse). Shown with a vertical red line is the median number of barcodes per element.

**Supplemental Figure 10.**
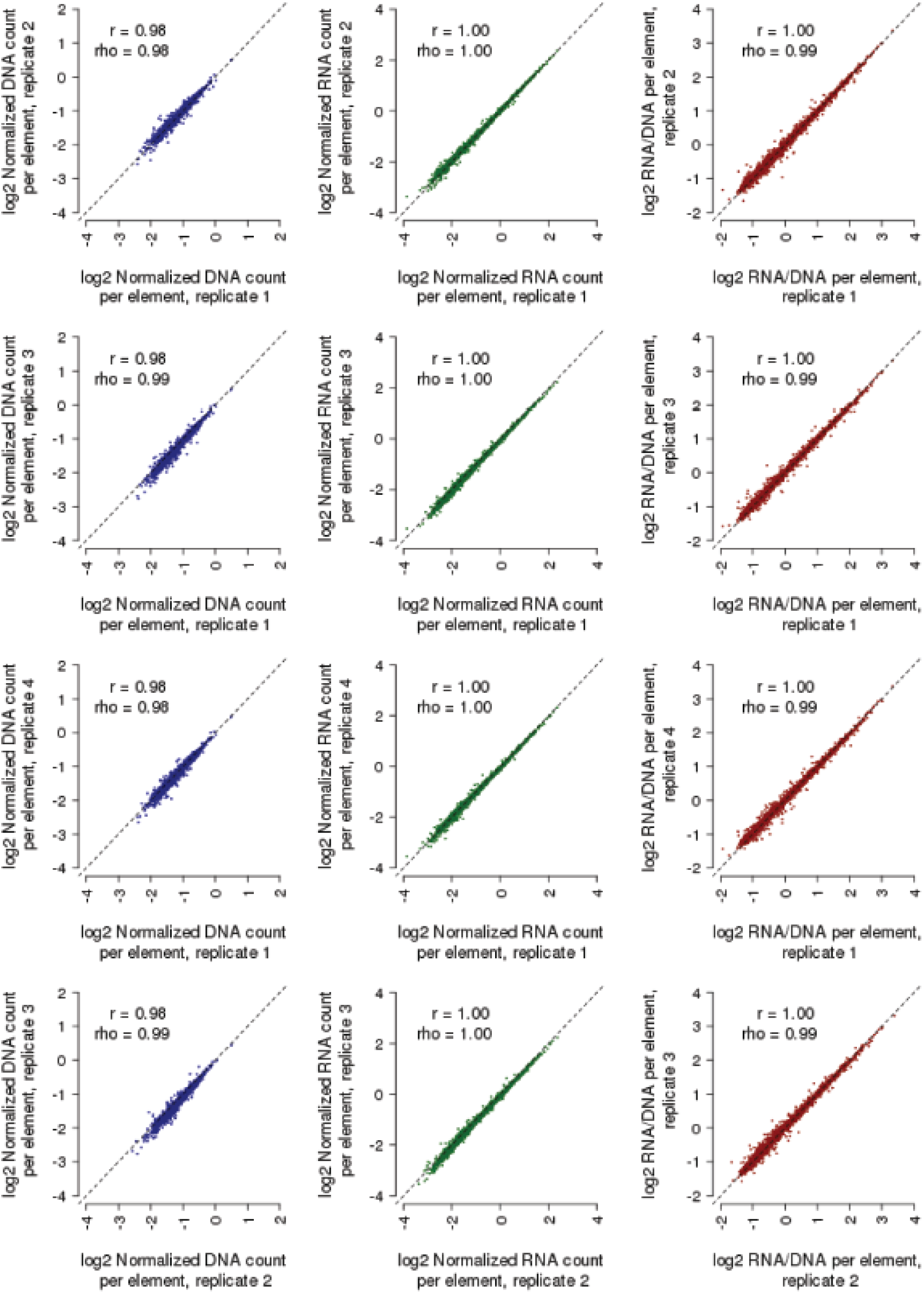
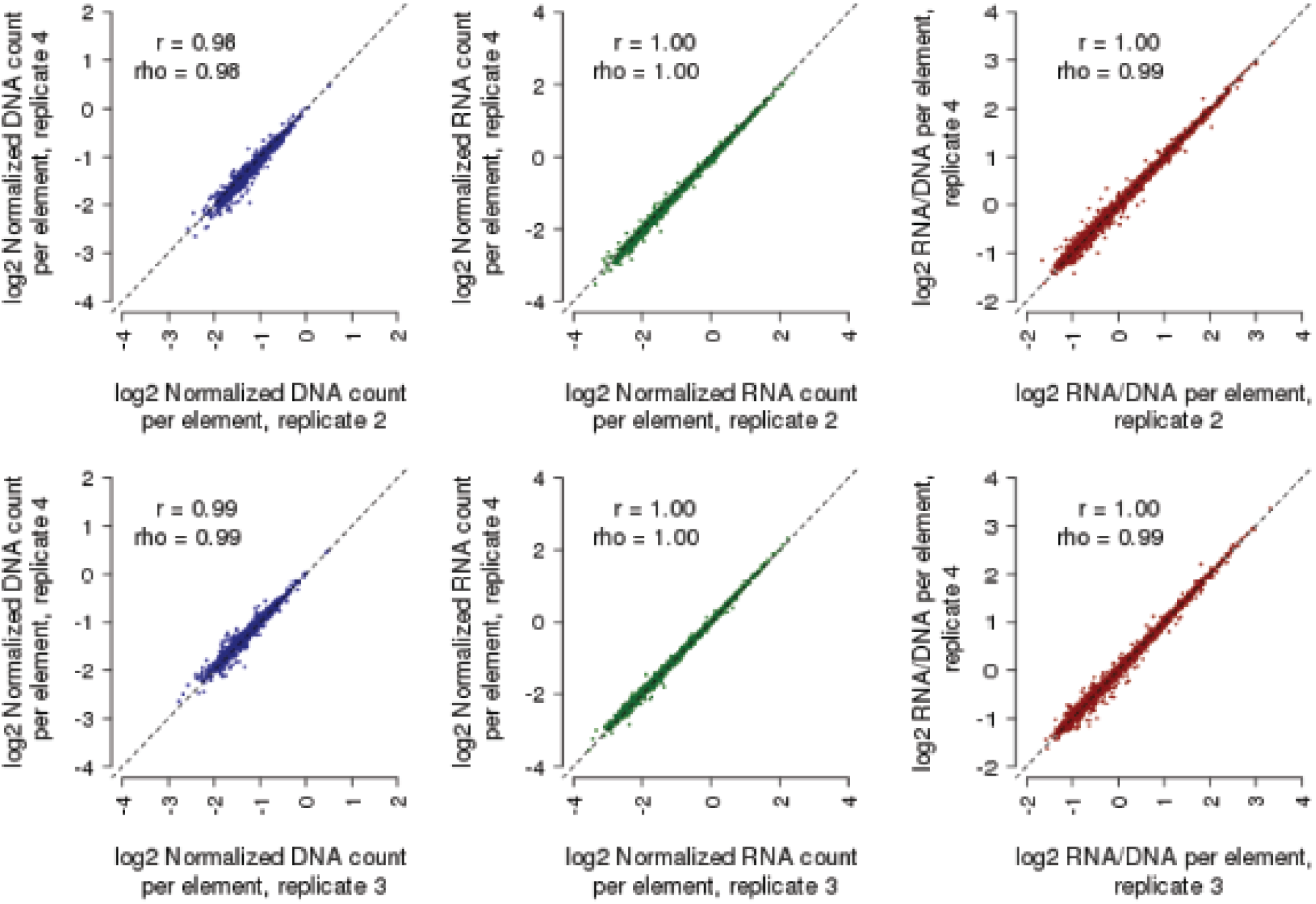
Reproducibility among replicates for the library testing orientation. Shown are scatter plots displaying the relationship between observed DNA counts (blue), RNA counts (green), and RNA/DNA ratios (red) for all pairwise comparisons among the four replicates, including data from both orientations (i.e., forward and reverse) tested. Also indicated are the Pearson (*r*) and Spearman (rho) correlation values. Candidate enhancers supported by fewer than 10 barcodes were filtered out prior to this analysis to reduce the impact of technical noise.

**Supplemental Figure 11.**
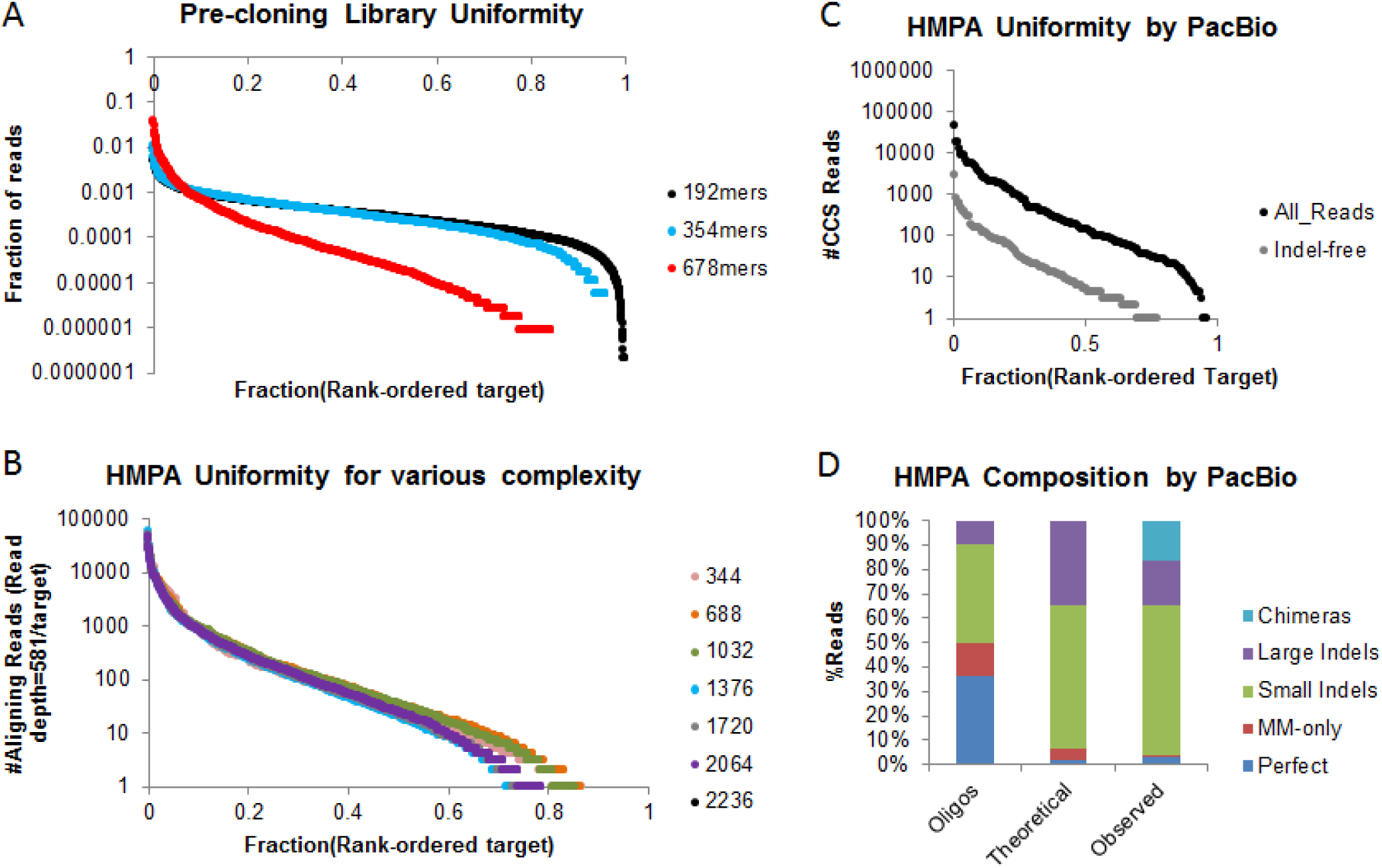
Evaluation of Multiplex Pairwise Assembly (MPA) and Hierarchical Multiplex Pairwise Assembly (HMPA) library quality. **A)** Plot of the uniformity of indel-free reads for 2,336 × 192mers (amplified off Agilent 230mer array), 2,336 × 354mers (after Multiplex Pairwise Assembly), and 2,236 × 678mers (after a single Hierarchical Multiplex Pairwise Assembly). The x-axis is rank-ordered according to the most to least abundant from left to right. The y-axis is the fraction of either indel-free reads (for 192mers and 354mers) or all reads (for 678ners). **B)** Uniformity for various HMPA reactions, with total number of target sequences ranging from 344 to 2,236. The sequencing reads were downsampled to normalize for the total number of targets (#Reads=581*#Targets). The Y axis is the number of downsampled reads from a given target sequence. **C)** One sub-library (of 172 targets), was sequenced on a PacBio Sequel. The plot shows the uniformity for all aligning reads (black) and indel-free reads (grey). **D)** Composition of the sub-library of 172 targets. The first column shows the breakdown of perfect, mismatch-only, small indels (<5 bp), large indels (≥ 5 bp), and chimeras, determined by Illumina sequencing. The second column shows the theoretical breakdown of 678mers based on each target consisting of four independent oligos. The third column shows the breakdown of 678mers based on PacBio sequencing.

**Supplemental Figure 12.**
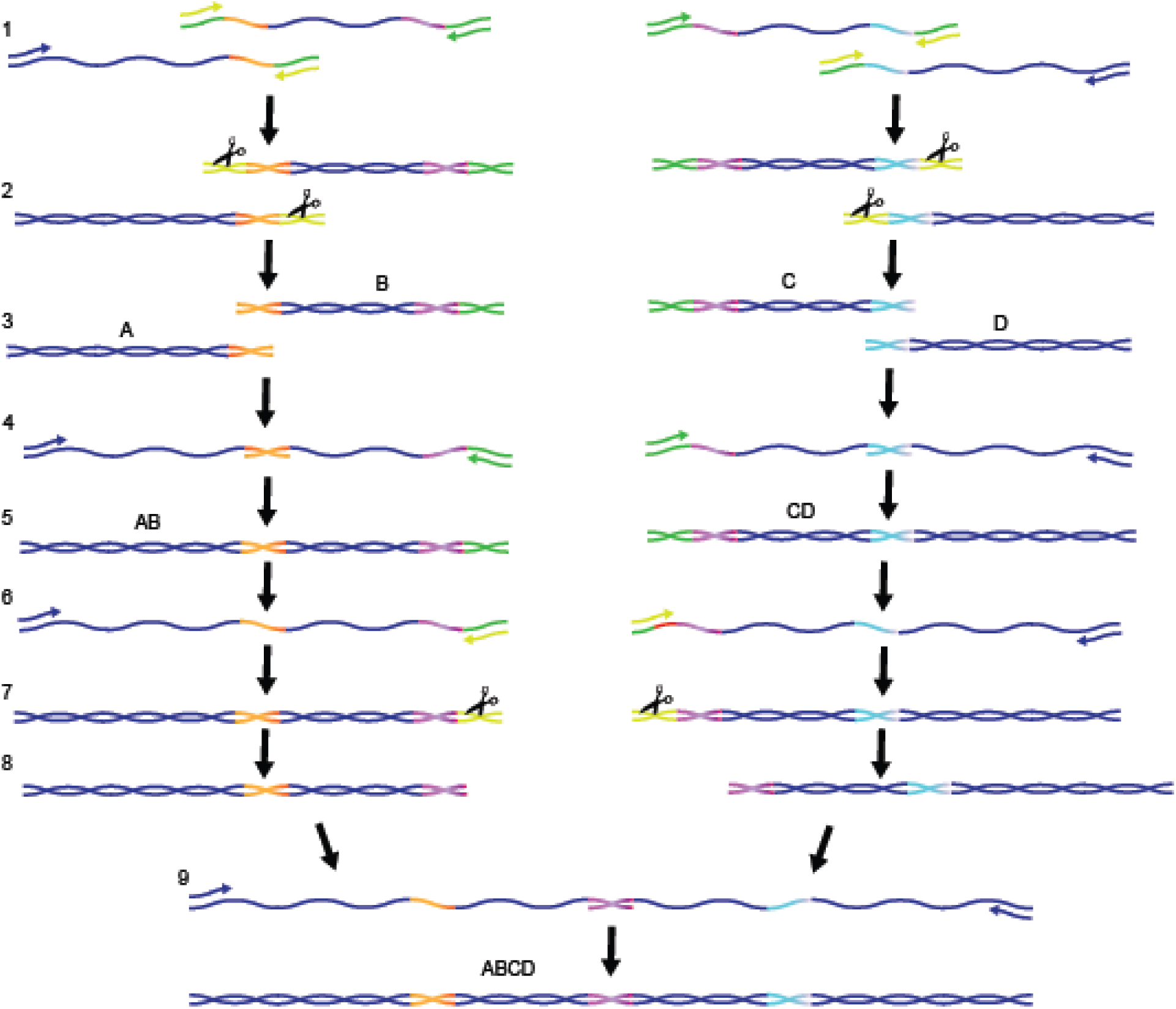
Hierarchical Multiplex Pairwise Assembly (HMPA) strategy. **1.** To generate a library of 678 bp enhancers, we ordered each enhancer as four oligonucleotides to be assembled (fragments “A”, “B”, “C”, and “D”). To assemble fragments “AB” and CD”, sequences were designed such that the 3’ end of fragments A and C had 30 bps of homology to the 5’ ends of fragments B and D, respectively (shown in red and orange). To remove adapter sequences from these ends of the fragments, uracil primers (shown in yellow) were used to incorporate uracils into the adapters during qPCR amplification. **2.** The resulting fragments were treated with USER enzyme (scissors) and put into an end-repair reaction, **3.** effectively removing the adapters. **4.** Fragments were assembled in a qPCR reaction by allowing the fragments to first anneal to one another for 5 cycles of PCR without primers, and then adding primers targeting the 5’ end of fragments A and C, and the 3’ ends of fragments B and D, **5.** resulting in fragments “AB” and “CD”. **6-9.** Adapter sequences were removed from the 3’ end of AB fragments and the 5’ end of CD fragments, and the final 678 bp “ABCD” enhancer sequences were assembled using the aforementioned assembly reaction.

**Supplemental Figure 13.**
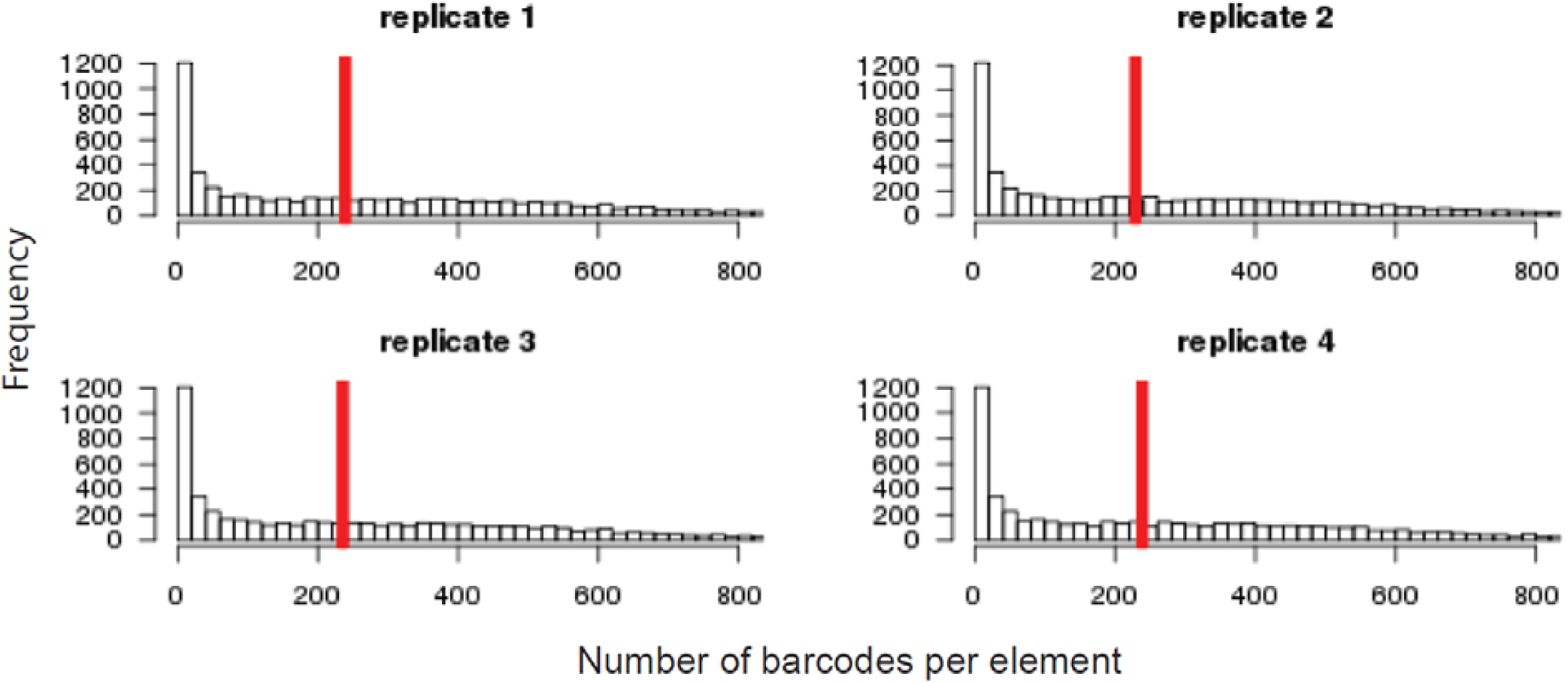
Histograms of barcodes per element for each replicate of the the library testing element size. Shown are histograms indicating the number of observed barcodes per element, for each of the 2,336 elements tested in each of the 4 replicates for the library testing size classes (i.e., short, medium, and long). Shown with a vertical red line is the median number of barcodes per element.

**Supplemental Figure 14.**
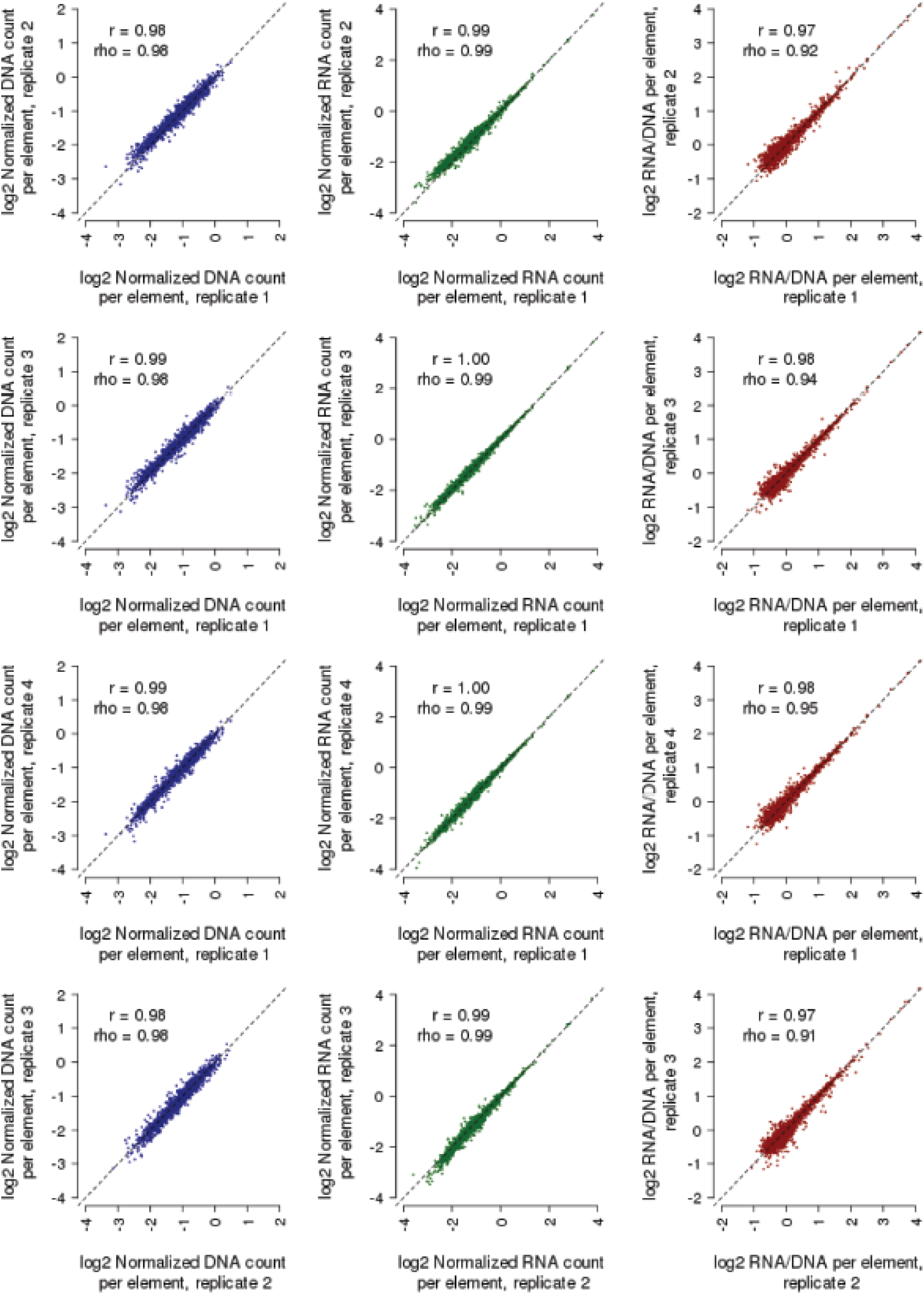
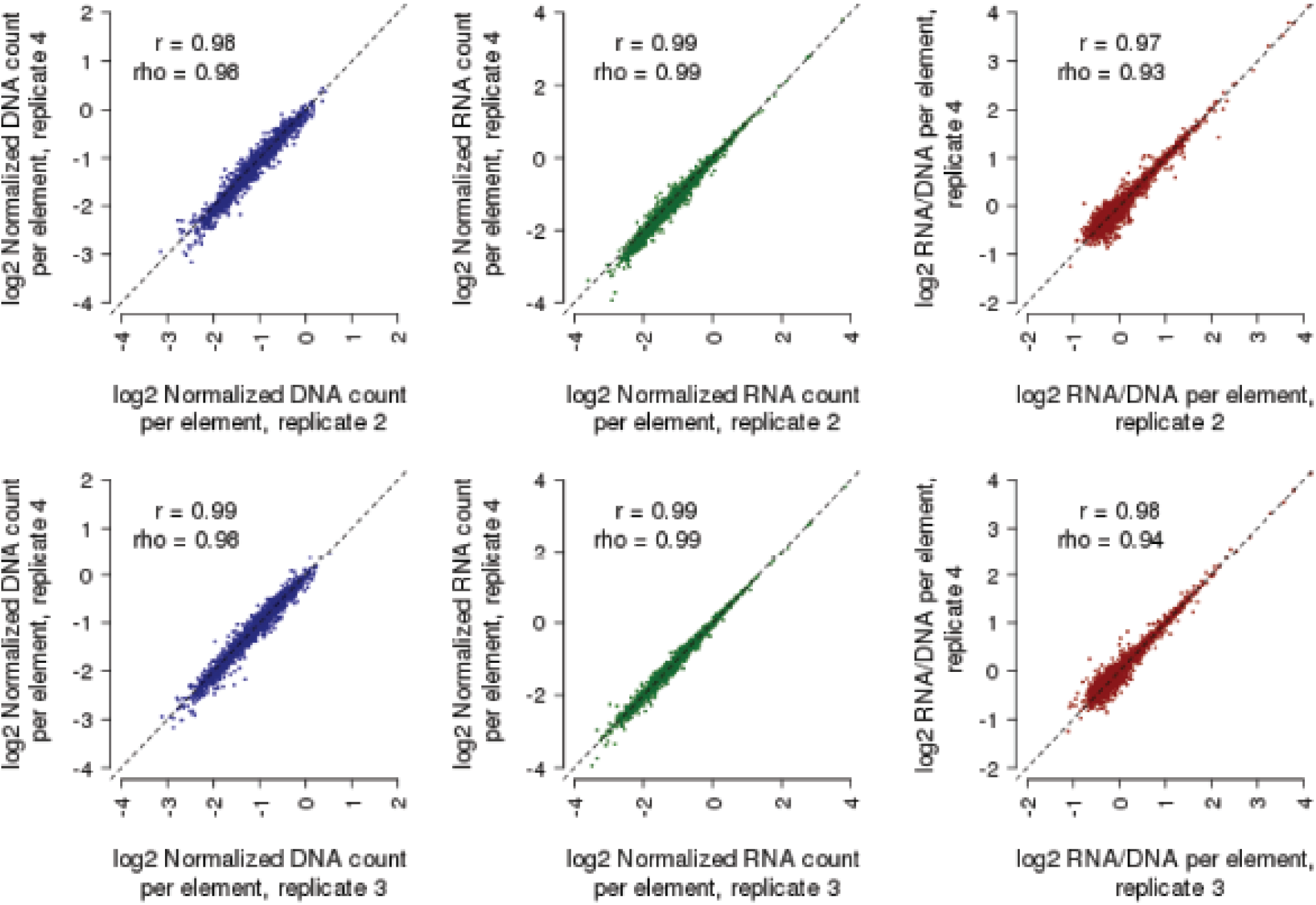
Reproducibility among replicates for testing of libraries of the same enhancers with different amounts of sequence context. Shown are scatter plots displaying the relationship between observed DNA counts (blue), RNA counts (green), and RNA/DNA ratios (red) for all pairwise comparisons among the four replicates, including data from all size classes (i.e., short, medium, and long) tested. Also indicated are the Pearson (*r*) and Spearman (rho) correlation values. Candidate enhancers supported by fewer than 10 barcodes were filtered out prior to this analysis to reduce the impact of technical noise.

**Supplemental Figure 15.**
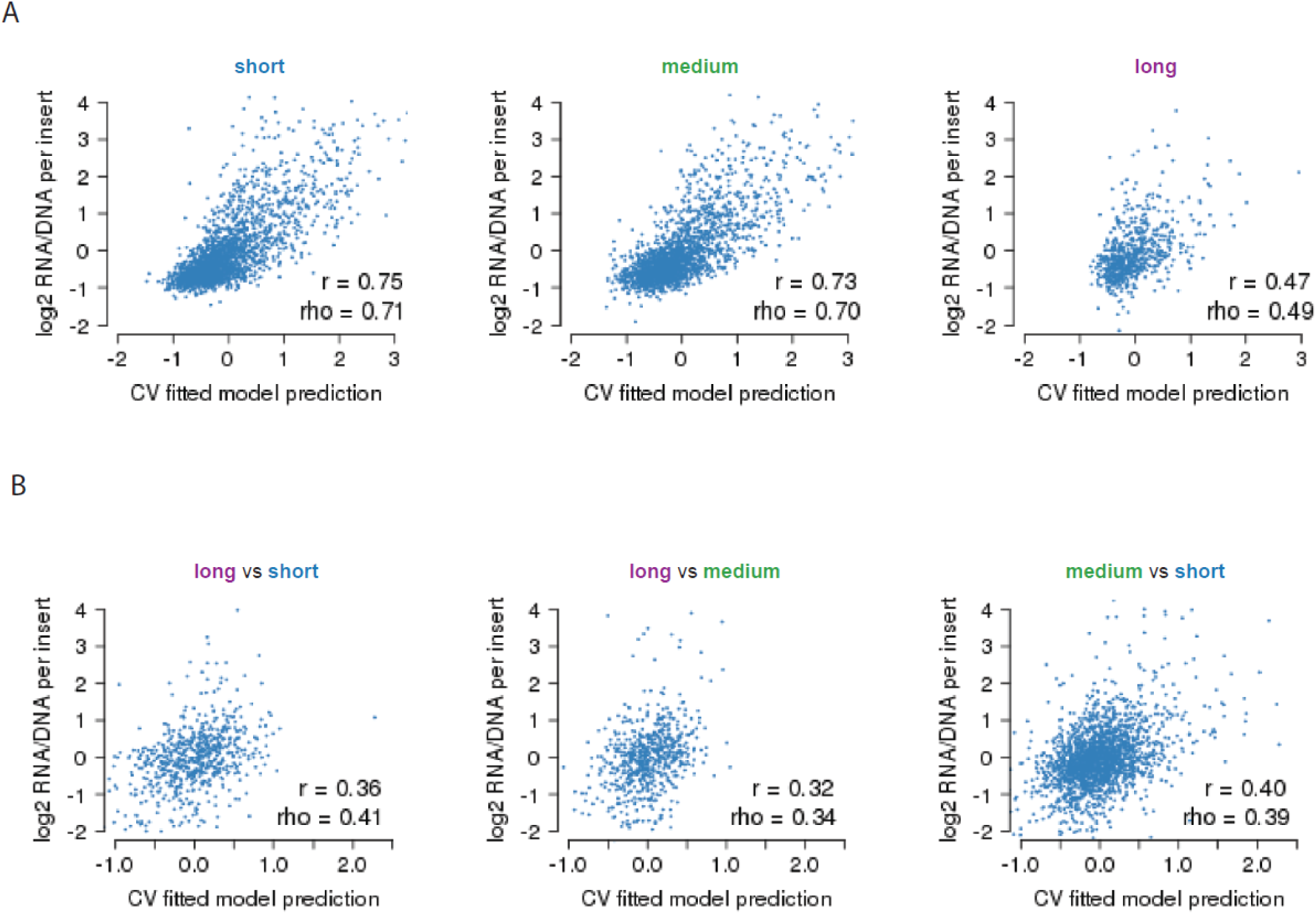
Performance of model-based predictions of the observed data from three size classes. **A-B)** Scatter plots displaying the relationship between the 10-fold cross-validated predictions derived from lasso regression models and the observed RNA/DNA ratios, for each of the 3 size classes tested (A) as well as the 3 corresponding differential comparisons tested (B). Also indicated are the Pearson (*r*) and Spearman (rho) correlation values.

## DATA AVAILABILITY

During the course of review, we intend to make a fully reproducible MPRA processing pipeline available at https://github.com/vagarwal87/MPRA-pipeline. The data will also be submitted to the Gene Expression Omnibus (GEO) for public use.

## AUTHOR CONTRIBUTIONS

J.K. and A.K. performed all cloning and sequencing for the 9 assays and all experimental work for orientation and length sections. J.K. and J.S. conceived of the HMPA protocol, and J.K. and A.K. developed and optimized it. A.K. produced schematic figures. V.A. performed the computational analyses, generated all remaining figures and tables, and adapted the MPRA analysis pipeline (which M.K. designed and contributed heavily towards) into a reproducible Github package. F.I. performed the transfections and lentiviral transductions for the 9 assays and wrote the associated methods sections. B.M. designed cloning steps and guided the development and testing of the MPRA assays. J.K., V.A., N.A., and J.S. wrote the remainder of the paper.

## ACKNOWLEDGEMENTS

We thank Seungsoo Kim and other members of the Shendure and Ahituv laboratories for general advice and critical feedback on the manuscript.

